# Characterising the Protein-Protein Interaction Between MDM2 and 14-3-3σ; Proof of Concept for Small Molecule Stabilisation

**DOI:** 10.1101/2023.09.26.559467

**Authors:** Jake A. Ward, Beatriz Romartinez-Alonso, Danielle F. Kay, Jeddidiah Bellamy-Carter, Bethany Thurairajah, Jaswir Basran, Hanna Kwon, Aneika C. Leney, Salvador Macip, Pietro Roversi, Frederick W. Muskett, Richard G. Doveston

## Abstract

Mouse Double Minute 2 (MDM2) is a key negative regulator of the tumour suppressor protein p53. MDM2 overexpression occurs in many types of cancer and results in the suppression of wild type p53. The 14-3-3 family of adaptor proteins are known to bind to MDM2 and the 14-3-3σ isoform controls MDM2 cellular localisation and stability to inhibit its activity. Therefore, small molecule stabilisation of the 14-3-3σ/MDM2 protein-protein interaction (PPI) is a potential therapeutic strategy for the treatment of cancer. In this work we provide a detailed biophysical and structural characterisation of the phosphorylation-dependent interaction between 14-3-3σ and peptides that mimic the 14-3-3 binding motifs within MDM2. The data show that di-phosphorylation of MDM2 at S166 and S186 is essential for high affinity 14-3-3 binding and that the binary complex formed involves one MDM2 di-phosphorylated peptide bound to a dimer of 14-3-3σ. Each of the two phosphorylated stretches of MDM2 occupies one of the two binding grooves of a 14-3-3σ dimer, a novel model for binding of di-phosphorylated peptides to 14-3-3 proteins. In addition, we show that the 14-3-3σ/MDM2 interaction is amenable to small molecule stabilisation. The natural product fusicoccin A forms a ternary complex with a 14-3-3σ dimer and an MDM2 di-phosphorylated peptide resulting in stablisation of the 14-3-3σ/MDM2 PPI. This work serves as a proof-of-concept of the drugability of the 14-3-3/MDM2 PPI and paves the way toward the development of more selective and efficacious small molecule stabilisers.

## Introduction

The transcription factor p53 plays a critical role in cell cycle regulation and tumour suppression.^1^ p53 inactivation via direct mutation or disruption of its regulatory network is a hallmark of many cancers.^2–4^ Therefore, p53 reactivation is a potentially powerful strategy for the development of antineoplastic drugs.^5^ In particular, the regulatory machinery that controls p53 homeostasis presents a number of targets for pharmaceutical intervention.^5,6^ One such target has been the protein-protein interaction (PPI) between p53 and its negative regulator MDM2 (Mouse Double Minute 2; MDM2 is used herein to refer to the human protein).^7^ MDM2 binds to the N-terminal transactivation domain of p53 to impede p53 interaction with DNA.^8^ In addition, MDM2 contains an E3 ubiquitin ligase domain that facilitates p53 ubiquitination and degradation via the proteasome.^9,10^ Direct inhibition of the p53/MDM2 PPI using small molecules is an effective approach for restoring p53 activity in malignant cells, but this has not yet yielded any clinically approved anti-cancer therapies.^5,11^ Targeting peripheral nodes of the p53 and/or MDM2 regulatory networks might provide a more tractable alternative or complimentary therapeutic strategy for p53 reactivation.

14-3-3 proteins represent one such node because they regulate MDM2 and p53 homeostasis via direct PPIs.^6^ 14-3-3 is a family of seven dimeric protein isoforms (β, γ, ε, ζ, η, σ and τ) that integrates and controls multiple signalling pathways.^12^ 14-3-3 proteins modulate the enzymatic activity, subcellular localisation, or interaction profile of over 1200 partner proteins via direct PPIs.^13,14^ 14-3-3 proteins typically recognise consensus phosphorylated motifs within disordered regions of partner proteins. Three types of phosphoserine (pS) or phosphothreonine (pT) peptide motifs are recognised by 14-3-3 proteins: mode I [RSX(pS/T)XP], mode II [RX(Y/F)X(PS/T)XP] or mode III [(pS/T)X-COOH].^15^ These peptide motifs bind to an amphipathic groove that is characteristic of 14-3-3 proteins, whereby the phosphate group interacts with a conserved basic binding pocket formed by residues K49, R56, R129 and R127 (14-3-3σ numbering).^12^ Although there is a high degree of sequence homology within the 14-3-3 family, each isoform exhibits its own affinity for the same binding partner,^16–18^ shows its own dimerisation behaviour,^19,20^ and exerts its own physiological response.^6,12^

For example, four 14-3-3 isoforms directly interact with the p53 C-terminus to positively regulate it in a manner dependent on Chk1/2 kinase activity.^21^ 14-3-3γ, ε, and ζ increase p53 transcriptional activity by promoting p53 tetramerisation and DNA binding, whereas 14-3-3σ achieves the same by increasing the half-life of p53 in the cell.^21^ Promoting 14-3-3/p53 PPIs using small molecules is, therefore, a potential therapeutic strategy for p53 reactivation and approaches for achieving this have been explored.^22,23^

14-3-3 proteins also regulate MDM2. 14-3-3σ binds to MDM2 via a phosphorylation-dependent interaction in addition to what is presumed to be a phosphorylation-independent interaction with the C-terminal RING domain of MDM2.^24^ This interaction negatively regulates MDM2 activity by promoting MDM2 auto-ubiquitination and degradation, and by sequestering MDM2 from the nucleus into the cytoplasm (Figure 1, panel A).^24^ This leads to stabilisation of p53 levels, and enhances p53 transcriptional activity.^24^ Thus, the 14-3-3σ/MDM2 PPI is also a potential therapeutic target for drug molecules that would stabilise the interaction and promote tumour suppression.

**Figure 1.**
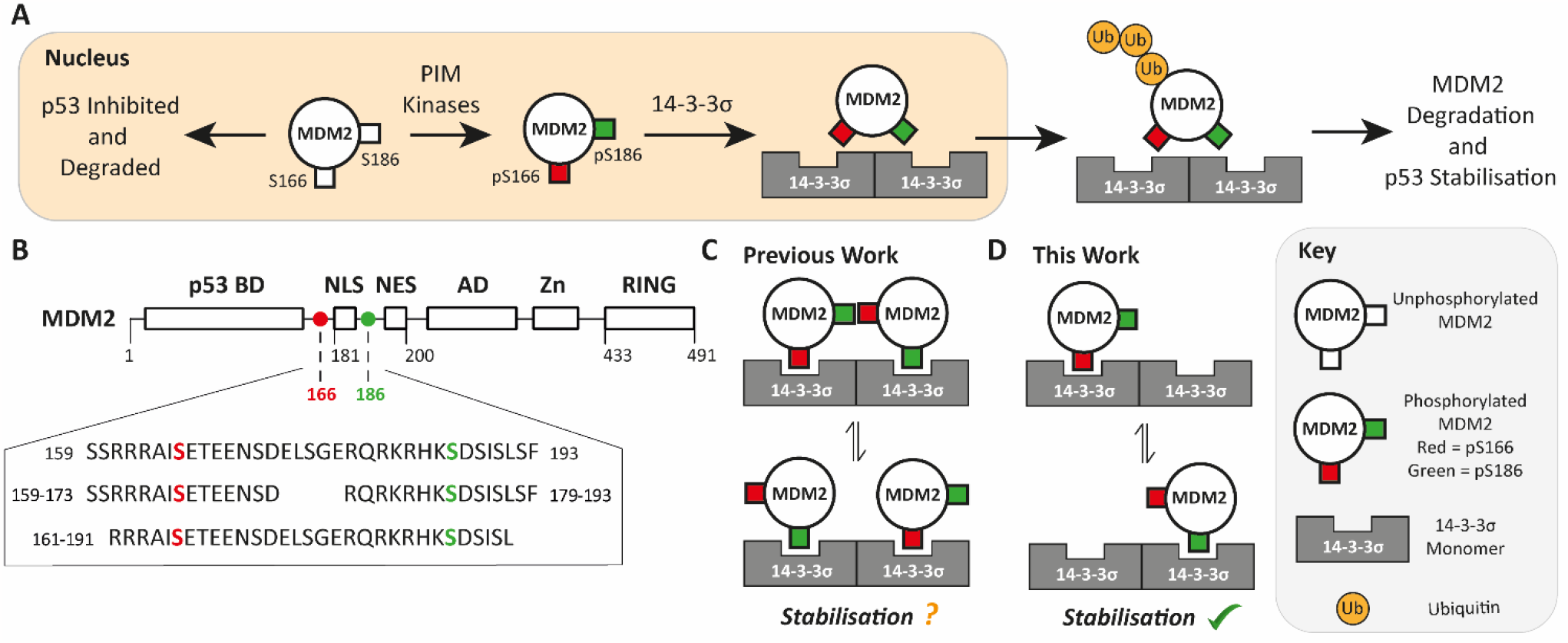
The role of 14-3-3σ in MDM2 regulation. **A**: 14-3-3σ interacts with phosphorylated MDM2 to promote its degradation and p53 stabilisation. **B**: MDM2 structure and 14-3-3 binding motifs. **C** and **D**: Proposed mechanistic models for the 14-3-3σ/MDM2 PPI.

The specific MDM2 residues and kinases that facilitate MDM2 binding to 14-3-3σ have not been identified through cellular investigations. The phosphorylated motifs that bind to 14-3-3 family proteins with highest probability, based on analysis of published examples using the *14-3-3Pred* webserver^25^ are sequences that incorporate pS166 and pS186 (Figure 1, panel B). This prediction is consistent with S166 and S186 phosphorylation by PIM kinases being a necessary condition for MDM2 binding to 14-3-3β, γ, ε, ζ, η, and τ isoforms.^26^ The physiological consequences of the interaction of MDM2 with these specific 14-3-3 isoforms have not been elucidated. However, in MDM2 the 14-3-3 binding motifs flank a nuclear localisation sequence (Figure 1, panel B). It is therefore likely that all 14-3-3 isoforms influence MDM2 cellular localisation, even though 14-3-3 isoform-specific profiles have not yet been characterised.

Fluorescence polarisation (FP) experiments recently provided the first *in vitro* evidence that all 14-3-3 isoforms, including 14-3-3σ, bind to peptides mimicking the pS166 and pS186 MDM2 motifs.^17^ The MDM2 peptides consistently bound to 14-3-3η with the highest affinity, and this isoform was used in further characterisation studies involving isothermal titration calorimetry (ITC) and surface plasmon resonance (SPR).^17^ 14-3-3σ showed the lowest affinity for the MDM2 pS166 and pS186 peptides. It is not uncommon for the σ isoform to form weaker interactions with phosphorylated peptides relative to the other isoforms.^16^ This might further point to the importance of the aforementioned phosphorylation-independent secondary interaction involving the MDM2 RING domain. FP showed that the pS166 MDM2 peptide bound all 14-3-3 isoforms with a higher affinity compared to the pS186 peptide.^17^ A crystal structure of 14-3-3σ in complex with the lower affinity pS186 MDM2 motif showed a peptide bound in each of the two binding grooves available in the 14-3-3 dimer with a mode I consensus binding pose.^17^ Significantly higher affinities for all 14-3-3 isoforms were observed using a MDM2 peptide with phosphorylation at both S166 and S186 sites.^17^ The higher affinity of di-phosphorylated peptides can be a result of the peptide simultaneously occupying both available binding grooves on a 14-3-3 dimer (i.e. 2:1 14-3-3 monomer to peptide stoichiometry). However, ITC data seemed to suggest that one peptide occupied each available 14-3-3 binding groove (i.e. 1:1 stoichiometry, Figure 1, panel C).^17^ Thus, it was proposed that high affinity binding of MDM2 for 14-3-3 proteins is driven by cooperativity between the two phosphorylation sites, a model described as ‘statistical rebinding’ (Figure 1, panel C).^17^

‘Multivalency’, where two or more phosphorylated motifs are required for high affinity binding, is integral to a number of 14-3-3 PPIs such as those with CFTR,^27^ LRRK2,^28^ tyrosine hydroxylase,^29^ IRSp53,^30^ MDMX,^17^ and MDM2.^17^ Such systems are usually heteroditopic and feature a ‘gatekeeper’ or ‘anchor’ phosphorylation site that binds 14-3-3 first, followed by the second site.^29,31^ The affinity of such interactions is dependent on the individual affinities of the two sites, and the effective molarity of the interacting partner.^31^ Such models assume that the two binding sites are identical, i.e. the 14-3-3 dimer does not exhibit any cooperative behaviour. It should be noted that the interactome profile of monomeric 14-3-3 protein differs from wild-type dimeric species indicating the physiological significance of the 14-3-3 dimer in partner protein recognition.^32^

In this manuscript we report on the further biophysical and structural characterisation of the interaction between 14-3-3σ and a pS166/pS186 MDM2 peptide using a combination of FP, ITC, native mass spectrometry (MS), protein X-ray crystallography and nuclear magnetic resonance (NMR). The data reveal that, in contrast to the previous report, the pS166/pS186 MDM2 peptide binds to 14-3-3σ via 2:1 stoichiometry, which is expected for a ‘bivalent’ interaction. The data also indicate that both phosphorylation sites do not simultaneously occupy each binding groove present in the 14-3-3 dimer. This points to a ‘rocking’ binding mechanism involving both phosphorylation sites (Figure 1, panel D). We also present data demonstrating that this interaction is amenable to small molecule stabilisation and therefore a viable drug target.

## Results and Discussion

Synthetic phospho-peptides were used to mimic the previously described 14-3-3 recognition motifs within the MDM2 sequence.^17,26^ MDM2 phosphorylated at S166 or S186 was mimicked by a mono-phosphorylated 15 amino acid peptide spanning residues 159-173 and 179-193 respectively: MDM2_159-173_^pS166^ and MDM2_179-193_^pS186^ (Figure 1, panel B). A di-phosphorylated 31 amino acid peptide (MDM2_161-191_^pS166/pS186^) was used to mimic MDM2 with phosphorylation at both sites (Figure 1, Panel B). The peptides were all amidated at the C-terminus, and either acetylated or fluorescently labelled at the N-terminus.

### MDM2 Di-Phosphorylation is Required for High Affinity Binding to 14-3-3σ

FP experiments were used to determine the relative binding affinities of the peptides for 14-3-3σ (Figure 2, panel A). 14-3-3σ was titrated to a fixed concentration of the fluorescently labelled peptides resulting in an increase in polarisation as the 14-3-3σ/MDM2 binary complex formed. The MDM2_159-173_^pS166^ peptide bound weakly to 14-3-3σ and saturation was not observed. As a result, an effective K_d_ could only be estimated to be > 115 µM. The MDM2_179-193pS186_ peptide showed stronger binding to 14-3-3σ, but again an accurate curve could not be fitted. An approximate K_d_ was estimated to be 26.2 ± 17.9 µM. In agreement with the previous study,^17^ the di-phosphorylated MDM2_161-191_^pS166/pS186^ peptide exhibited significantly higher affinity binding to 14-3-3σ (effective K_d_ = 2.9 ± 1.0 µM).

**Figure 2.**
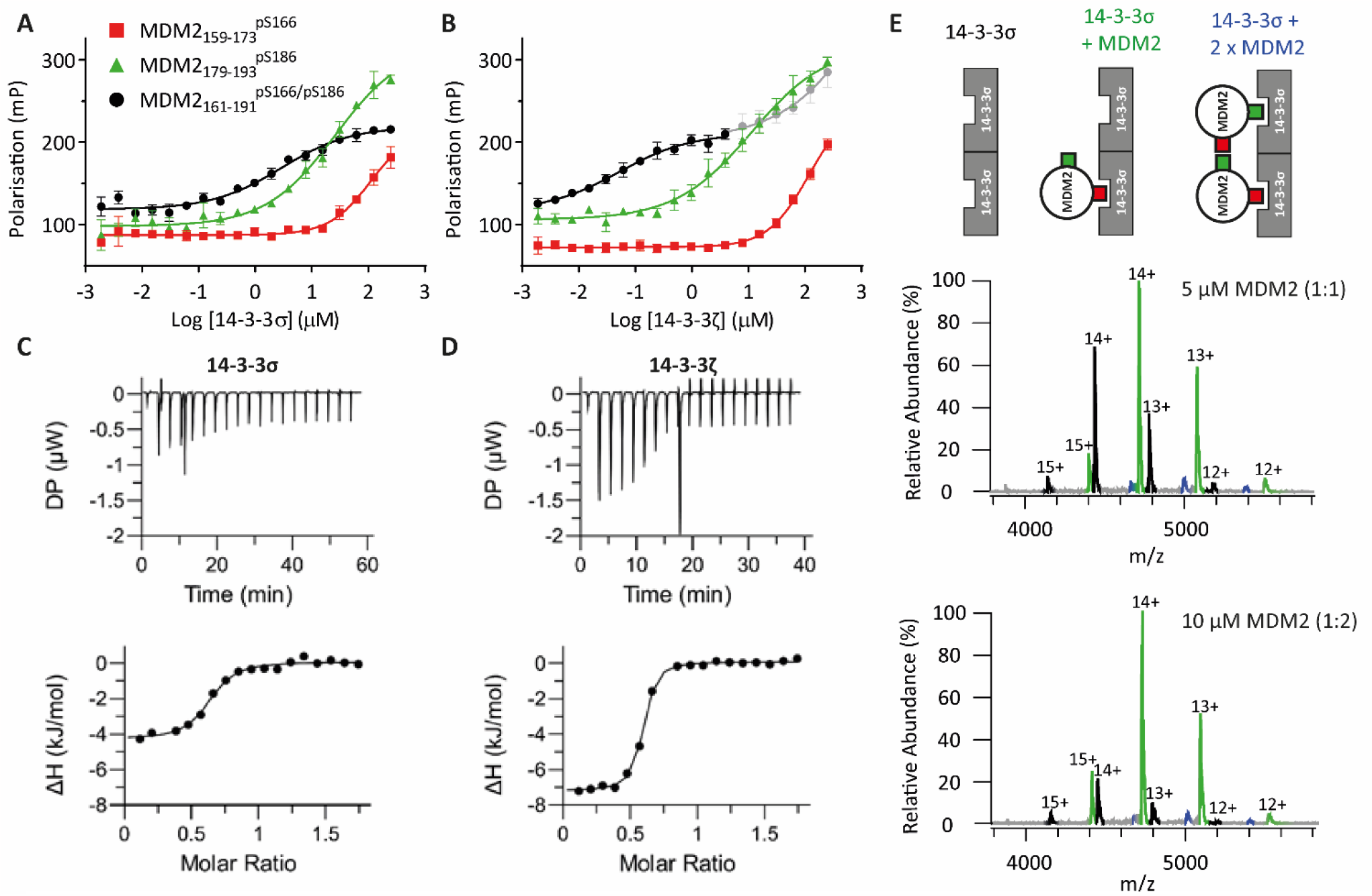
Biophysical investigation of the 14-3-3σ/MDM2 PPI. **A-B**: FP data for MDM2 peptide binding to 14-3-3σ (**A**) and 14-3-3ζ (**B**, the second binding event is shown in grey). 14-3-3 protein was titrated to 10 nM FITC-labelled peptide in buffer containing 25 mM HEPES pH 7.5, 100 mM NaCl, 10 mM MgCl_2_, 0.1% v/v Tween20, 0.1 mg/mL BSA and 1 % v/v DMSO. Error bars represent SD for n=3 replicates. **C-D**: ITC data for MDM2 peptide binding to 14-3-3σ (**C**) and 14-3-3ζ (**D**). Acetylated MDM2_161-191_^pS166/pS186^ peptide (1.0 mM) was titrated to 14-3-3 protein (0.1 mM) at 25 °C in buffer containing 25 mM HEPES pH 7.5, 100 mM NaCl, 10 mM MgCl_2_ and 1 % v/v DMSO. The TFA content of the peptide was determined by ^19^F-NMR in order to accurately calculate peptide concentration (see Fig. S5 and SI for details). **E**: Native mass spectrometry data confirming 2:1 stoichiometry of 14-3-3σ monomer to MDM2 peptide. All protein concentrations are given as 14-3-3 monomer concentrations

The maximum polarisation level observed for the MDM2_161-191_^pS166/pS186^ peptide was lower than that for the MDM2_179-193_^pS186^ peptide (Figure 2, panel A). This suggests that the N-terminal fluorophore of the MDM2_161-191_^pS166/pS186^ peptide peptide has greater rotational flexibility when bound to 14-3-3σ compared to the shorter MDM2_179-193_^pS186^ peptide. This might suggest that the C-terminal pS186 site indeed has a higher affinity for 14-3-3 compared to the pS166 site as indicated by the initial FP data.

In the previous study it was shown that other 14-3-3 isoforms, e.g. 14-3-3ζ, bound with higher affinity that 14-3-3σ to phosphorylated MDM2 peptide motifs.^17^ To corroborate this, 14-3-3ζ was titrated to fixed concentrations of the three peptides (Figure 2, panel B). As with 14-3-3σ, the MDM2_159-173_^pS166^ peptide bound weakly to 14-3-3ζ, and no effective K_d_ value could be obtained. The MDM2_179-193_^pS186^ peptide bound to 14-3-3ζ with approximately a 2-fold higher affinity compared to the σ isoform (effective 14-3-3ζ K_d_ = 11.36 ± 5.4 µM vs 14-3-3σ K_d_ = 26.2 ± 17.9 µM). The MDM2_161-191_^pS166/pS186^ peptide again bound with significantly higher affinity, and in line with the previous study, a biphasic curve was observed.^17^ Biphasic binding curves for 14-3-3 PPIs have only been reported for MDM2 and MDMX peptides through FP studies.^17^ The observation is difficult to rationalise because FP only reports on the global average of all binding events that take place with a significant change in mass, hence ‘effective’ K_d_. Thus, the biphasic curve cannot be related to peptide-protein binding events. It could be related to non-specific reduction of fluorophore rotational freedom, or higher order complex formation at high protein concentrations, but further investigation is required. Nevertheless, assuming the monophasic binding curve observed for 14-3-3σ reflects the overall weaker affinity of this isoform for the MDM2_161-191_^pS166/pS186^ peptide, the effective K_d_ for 14-3-3ζ in the first phase was 56-fold lower than that for 14-3-3σ (51.9 ± 28.3 nM vs. 2.9 ± 1.0 µM). The effective K_d_ for the second phase could not be obtained because the curve did not reach saturation.

Further FP experiments were conducted to confirm that di-phosphorylation was a requirement for high affinity of the MDM2_161-191_^pS166/pS186^ peptide to 14-3-3σ. 14-3-3σ was titrated to 35 amino acid peptides with single phosphorylation sites at pS166 and pS186: MDM2_159-193_^pS166^ and MDM2_159-193_^pS186^. The longer MDM2_159-193_^pS166^ peptide showed a small increase in the maximum polarisation value obtained upon 14-3-3σ titration compared to the 15 amino acid MDM2_159-173_^pS166^ peptide (Figure S2). As before, saturation was not observed and an effective K_d_ value could not be obtained. The longer MDM2_159-193_^pS186^ peptide bound to 14-3-3σ with much weaker affinity compared to the 15 amino acid MDM2_179-193_^pS186^ peptide. Saturation was not observed and an effective Kd value could only be estimated to be > 124 µM (Figure S2). Therefore, di-phosphorylation and not peptide length, accounted for the higher affinity of the MDM2_161-191_^pS166/pS186^ peptide. A noteworthy observation was that the maximum polarisation level observed for the longer MDM2_159-193_^pS186^ peptide was comparable to that observed for the di-phosphorylated MDM2_161-191_^pS166/pS186^ peptide. In contrast, the shorter MDM2_179-193_^pS186^ peptide exhibited a much higher maximum polarisation value (*vida supra*). These data further point to increased rotational flexibility of the N-terminal fluorophore when pS186 but not pS166 is anchored in the 14-3-3σ phosphate binding pocket.

To confirm that the MDM2_161-191_^pS166/pS186^ peptide was engaging in a specific interaction with the 14-3-3σ binding groove, competition FP experiments were performed. The N-terminally acetylated analogue of the MDM2_161-191_^pS166/pS186^ peptide was titrated to a fixed concentration of fluorescently labelled MDM2_161-191_^pS166/pS186^ and 14-3-3σ. As the concentration of the unlabelled peptide increased, polarisation decreased because the fluorescent tracer peptide was competed out of the binding site (Figure S3). The unlabelled MDM2_161-191_^pS166/pS186^ peptide had an IC_50_ of 3.2 ± 0.5 µM. A peptide mimicking the mode III 14-3-3 binding motif of the ERα transcription factor was also investigated. The 14-3-3/ERα PPI has been fully characterised, and this peptide motif is known to engage with the 14-3-3σ binding groove.^33^ This peptide also competed for binding to 14-3-3σ with the fluorescently labelled MDM2_161-191_^pS166/pS186^ tracer peptide with an IC_50_ of 1.5 ± 0.2 µM (Figure S3). Thus, it can be concluded that the MDM2_161-191_^pS166/pS186^ peptide binds to the 14-3-3σ binding groove in a manner that depends on 14-3-3 recognition of MDM2 with phosphorylation at S166 and S186.

ITC provided orthogonal conformation of the K_d_ values obtained, and further insight into the thermodynamics and stoichiometry of the MDM2 peptide interactions with 14-3-3σ and ζ. In these experiments N-acetylated MDM2 peptides were titrated to a fixed concentration of 14-3-3 protein. Binding of the mono-phosphorylated MDM2_159-173_^pS166^ and MDM2_179-193_^pS186^ peptides could not be detected (Figure S4). However, ITC could detect MDM2_161-191_^pS166/pS186^ peptide binding to 14-3-3σ with a K_d_ of 1.5 ± 0.4 µM (Figure 2, panel C), and to 14-3-3ζ with ~4-fold higher affinity (K_d_ = 0.38 ± 0.053 µM, Figure 2, panel D), values comparable with the FP data (Table 1). In these experiments biphasic binding was not observed for 14-3-3ζ. This could be due to the different concentration regimes used in ITC vs. FP, or could be due to the absence of the fluorophore in the ITC experiments. The higher affinity of the peptide for 14-3-3ζ was a result of a more negative enthalpy contribution relative to 14-3-3σ. Overall however, the interaction of the MDM2_161-191_^pS166/pS186^ peptide with both 14-3-3 isoforms was entropically driven (Table 1). This is interesting because the interactions of di-phosphorylated peptides that span the 14-3-3 dimer and simultaneously engage both binding grooves are typically enthalpy-dominated.^34^ The data seem to suggest that the MDM2 peptide binds to the 14-3-3 dimer in more than one way, and/or that MDM2 peptide binding displaces previously ordered water molecules in the binding groove.

**Table 1.**
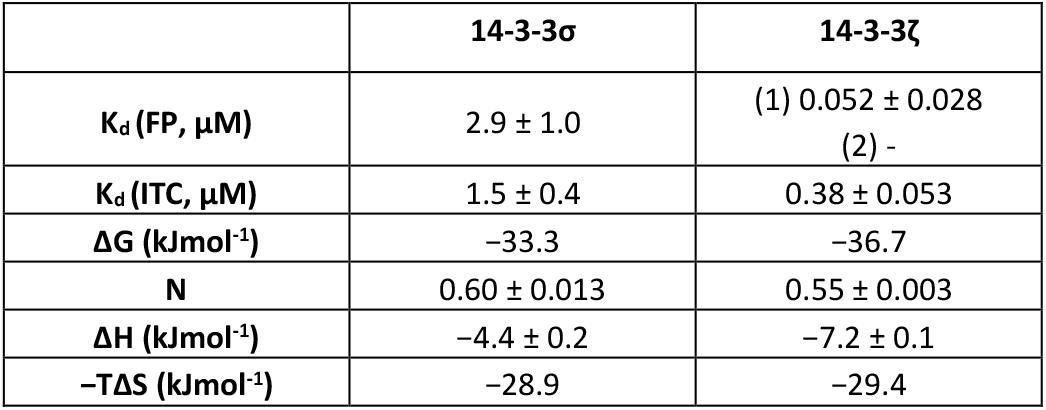
K_d_ values, thermodynamic and kinetic parameters for MDM2_161-191_^pS166/pS186^ peptide binding to 14-3-3σ and 14-3-3ζ.

### One MDM2 Di-Phosphorylated Peptide Binds to a 14-3-3 Dimer

The ITC experiments showed that the stoichiometry (N) for the interactions of the MDM2_161-191_^pS166/pS186^ peptide with 14-3-3σ and 14-3-3ζ to be 0.60 and 0.55 respectively (Table 1, Figure 2 panels C-D).^§^ This is a strong indication that the 14-3-3 dimer interacts with a single di-phosphorylated MDM2_161-191_^pS166/pS186^ peptide, i.e. 2:1 ratio of 14-3-3 monomer to peptide. These data disagreed with that from the previous study which reported 1:1 binding stoichiometry.^17^ To confirm the ITC data native MS experiments were conducted. Native MS allows for the analysis of proteins and protein complexes without denaturing protein secondary structure or disrupting non-covalent interactions.^35^ Stoichiometry and binding equilibria can be determined from the unique mass of the complexes involved.^36^ MS analysis revealed that 14-3-3σ was almost exclusively dimeric at a concentration of 5 µM (Figure S7).

Upon the addition of an equimolar concentration of the MDM2_161-191_^pS166/pS186^ peptide a binary complex was observed (Figure 2, panel E). The binary complex consisted of the 14-3-3σ dimer and a single MDM2_161-191_^pS166/pS186^ peptide. When a higher concentration of the MDM2_161-191_^pS166/pS186^ peptide was added the population of the binary complex increased, and the population of apo-14-3-3σ decreased (Figure 2, panel E). There was scant evidence of a complex consisting of a 14-3-3σ dimer and two MDM2_161-191_^pS166/pS186^ peptides, and that which was observed is most likely a result of non-specific binding.^37^ Therefore, the native MS data confirm that the interaction of 14-3-3σ with the MDM2_161-191_^pS166/pS186^ peptide has a stoichiometry of 2:1. This contrasts to native MS analysis of other 14-3-3 PPIs which involve 1:1 binding of phospho-peptide to 14-3-3 monomer.^36^

### Structural Analysis of the 14-3-3σ/MDM2 PPI

A crystal structure of the MDM2_161-191_^pS166/pS186^ peptide in complex with a 14-3-3σ ΔC construct (lacking the final 17 residues at the C-terminus) was determined to a resolution of 1.3 Å. The complex of the 14-3-3σ dimer shows density for a single peptide occupying both 14-3-3σ binding grooves (Figure 3, panels A and B). In the crystal the asymmetric complex sits at the same site (around a crystal twofold axis) in two different orientations, so that the ordered parts of the N-terminal (residues ^164^AIpSE^167^) and C-terminal (^184^HKpSD^187^) portions of the peptide overlap (static disorder, Figure 3, panels A and B). The peptide binds in the expected orientation with its C-terminal end directed toward the N-terminus of 14-3-3σ. Both pS166 and pS186 phosphorylation sites are engaged with the 14-3-3σ binding groove, interacting with the residues that form the phosphate binding pocket: R56, R129 and Y130. The -2 and +1 residues relative to either pS are also well ordered in the crystal (Figure 3, panel C-D).

**Figure 3.**
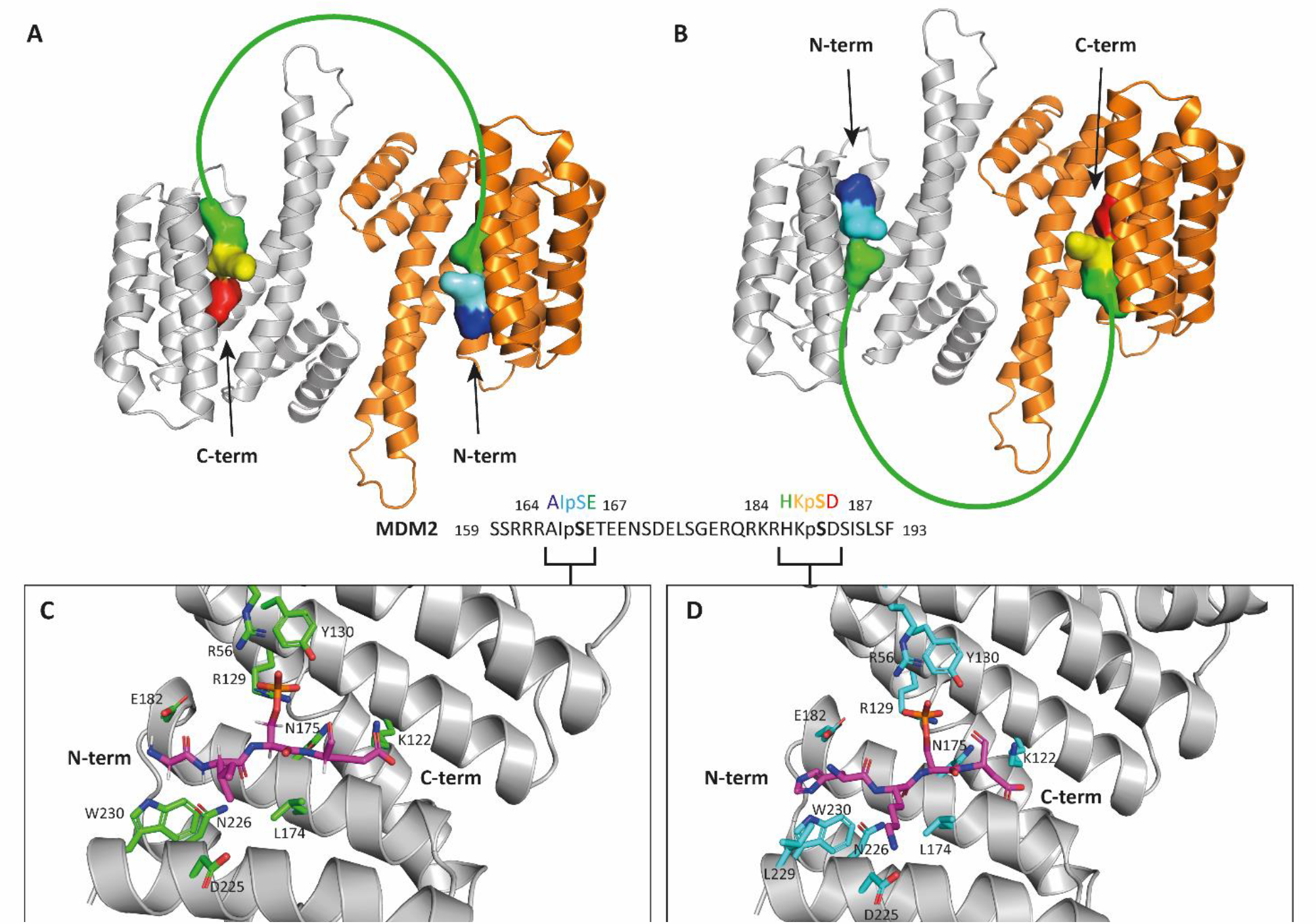
Crystallographic analysis of the 14-3-3 interaction with the MDM2_161-191_^pS166/pS186^ peptide (PDB: 8P0D). **A-B**: The 14-3-3σ dimer in the crystal, in cartoon representation, with one monomer shown in grey and one monomer in orange. The complex sits in the same site in two different orientations (**A** and **B**) causing the spatial overlap of the two ordered regions of the MDM2 peptide in the crystal. The latter are represented as surfaces in panels **A** and **B**. The 164-167 stretch of the MDM2 peptide is coloured blue to green from N-terminus to C-terminus; the 184-187 stretch of the MDM2 peptide is coloured green to red from N-terminus to C-terminus. The disordered part of the MDM2 peptide, 168-183 is represented as a green line. **C-D**: The two overlapping ordered stretches of the MDM2 peptide bound in the 14-3-3σ binding groove. N atoms are shown in blue, O atoms in red. Peptide: in sticks with C atoms in magenta. 14-3-3: backbone in grey cartoon representation; side chains that interact with the peptide in sticks representation with C atoms in green or cyan. **C**: the MDM2 164-167 stretch. **D**: the MDM2 184-187 stretch.

MDM2 peptide main chain atoms interact with the side chains of 14-3-3σ N175 and N226 and, together with the pS side chain interactions, define the pose of the peptide backbone around the phosphorylation site in a MDM2 sequence-independent manner. Polar interactions are observed between main chain O and N atoms of MDM2 D157 and E167 and the side chain of 14-3-3σ N175; and between main chain O and N atoms of MDM2 I165 and K185 and the side chain of 14-3-3σ N226.

Further to these MDM2 peptide backbone interactions there are some MDM2 peptide sequence-dependent side chain interactions. In the ^164^AIpSE^167^ part of the peptide, a hydrophobic contact is observed between the side chains of MDM2 I165 and 14-3-3σ L229 whilst MDM2 E167 forms a salt bridge with 14-3-3σ K122. H-bonds are observed between the side chains of MDM2 H184 and 14-3-3σ Y181, E182 and W230; and between the side chains of MDM2 K185 and 14-3-3σ D225; MDM2 D187 forms a salt bridge with 14-3-3σ K122. The interactions of the ^184^HKpSD^187^ part of the peptide, confirm the ones already observed in PDB ID 6YR6^17^ and more broadly both peptides bind in a similar way to other 14-3-3 client peptides (see Figure S9). The structure was deposited with PDB ID 8P0D.

Protein NMR was used to further investigate the stoichiometry and dynamics of the 14-3-3σ/MDM2 PPI. Backbone chemical shift assignments (^15^N, ^13^C, ^1^H) have previously been reported for the 14-3-3σ ΔC construct,^38^ and the C-terminal domain of 14-3-3σ.^39^ This enabled ^1^H-^15^N transverse relaxation optimised spectroscopy (TROSY) spectra to be collected for ^15^N-enriched 14-3-3σ (full length), and the observed resonances were assigned to specific amino acid residues. Only a single resonance is observed for each amide proton/nitrogen pair in the spectrum indicating that 14-3-3σ is observed as a symmetric dimer in solution (although two distinct complexes could be distinguished by crystallography, the system is under fast exchange, and the two complexes would be equivalent in solution).

^1^H-^15^N TROSY spectra were collected on 14-3-3σ in complex with the MDM2_161-191_^pS166/pS186^ peptide and compared to the spectrum of free 14-3-3σ. In the presence of 2 molar equivalents of peptide relative to 14-3-3σ monomer distinct chemical shift perturbations (CSP) were observed (Figure 4, panel A, B). There were no differences in the CSPs observed when 2.5 molar equivalents of peptide relative to 14-3-3σ monomer were used (Figure S10). This corroborates the ITC data that indicates binding saturation at two molar equivalents of peptide. It provides further evidence for 2:1 14-3-3σ monomer-peptide binding stoichiometry because a greater molar excess of peptide would be expected to be required to reach saturation if two peptides were simultaneously engaging a 14-3-3σ dimer. The symmetry of the system was however retained upon peptide binding as shown by the single distinct set of resonances observed. This suggests that the MDM2_161-191_^pS166/pS186^ peptide engages with both monomers via a fast exchange mechanism. CSPs were mapped onto the crystal structure of 14-3-3σ bound to the MDM2_161-191_^pS166/pS186^ peptide (Figure 4, panel B). CSPs were observed for four amino acids within, or proximal to, the 14-3-3σ binding groove: S45, V134, F176 and S186. The most significant perturbation was observed for S45 which lies at the N-terminal end of the binding groove and the crystal structure indicates that S45 forms long hydrogen bonds with the carboxylate side chains MDM2 E167 and E187. V134 lies at the C-terminal entrance to the binding groove and faces the main 14-3-3 dimer interface. Thus, it is possible the CSP results from the MDM2 peptide spanning the length of the binding groove, and perhaps distortion of the dimer upon binding. F176 lies next to N175 which is observed to form a polar contact with the MDM2 peptide backbone in the crystal structure. S186 also lies at the C-terminal entrance to the binding groove and is observed to point away from the peptide in the crystal structure, perhaps indicating this residue side chain rotates upon peptide binding.

**Figure 4.**
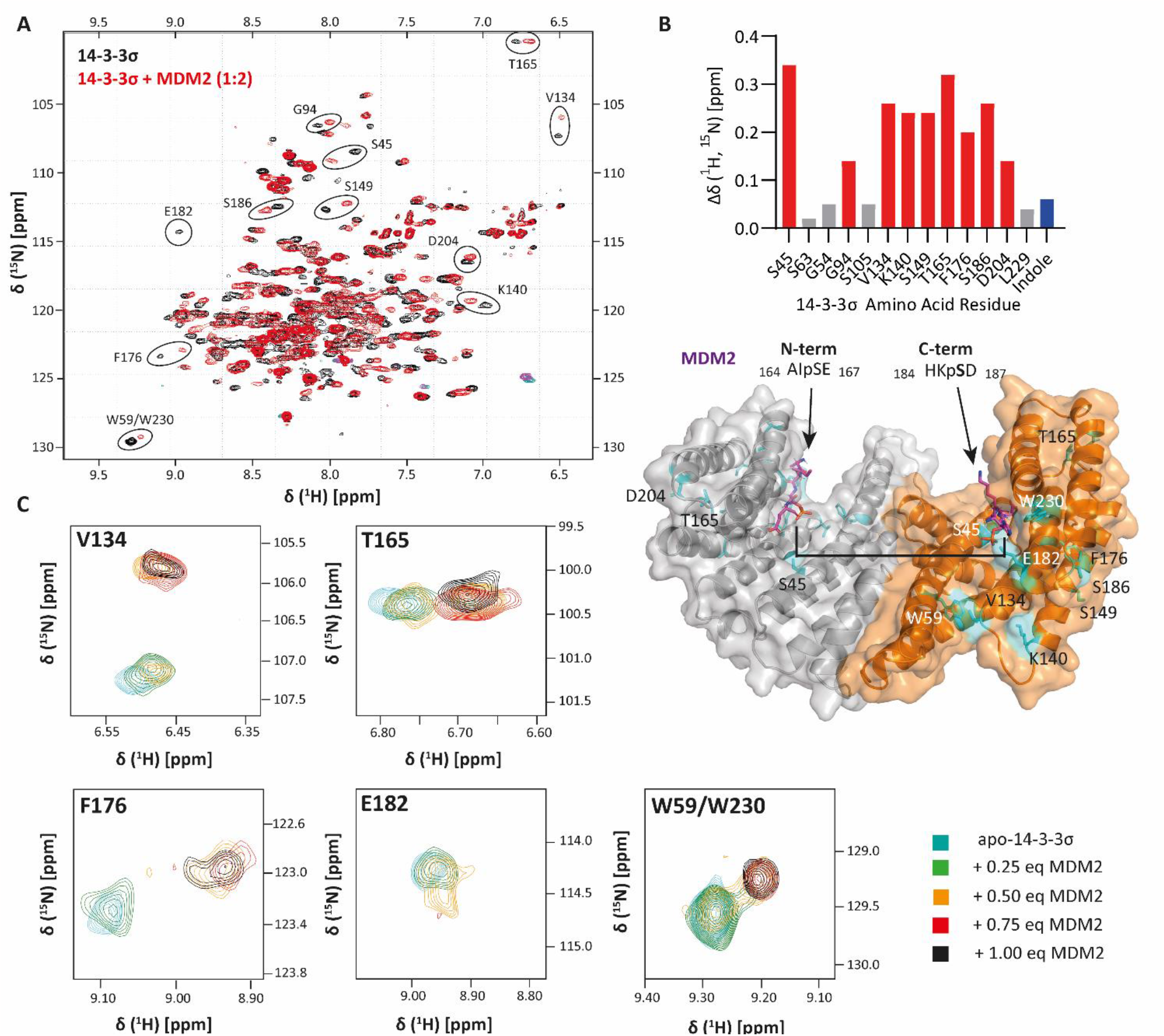
^1^H-^15^N TROSY NMR analysis of the 14-3-3σ interaction with the MDM2_161-191_^pS166/pS186^ peptide. **A**: Superimposed ^1^H-^15^N TROSY spectra of ^15^N-labelled apo-14-3-3σ (100 µM, black) and ^15^N-labelled 14-3-3σ (100 µM) in complex with the MDM2_161-191_^pS166/pS186^ peptide (200 µM) (red). All protein concentrations are given as 14-3-3 monomer concentrations. **B, top:** Plot showing the notable combined ^1^H and ^15^N chemical shift perturbations in the spectrum of ^15^N-labelled 14-3-3σ (100 µM) in complex with the MDM2_161-191_^pS166/pS186^ peptide (200 µM). Residues displaying significant CSPs are shown in red, the CSP relating to the indole resonance is shown in blue. For reference the CSP for three residues that did not show notable changes are shown in grey. **B, bottom**: CSPs (cyan) mapped onto the protein X-ray crystal structure of 14-3-3σ ΔC dimer in complex with the MDM2_161-191_^pS166/pS186^ peptide (purple). The disordered part of the MDM2 peptide, 168-183 is represented as a black line **C**: Selected ^1^H-^15^N TROSY resonances for 14-3-3σ (100 µM) in the presence of increasing concentrations of the MDM2_161-191_^pS166/pS186^ peptide (see key).

There were notable CSPs for two resonances within the indole region of the spectra. 14-3-3σ contains two tryptophan residues bearing indole side chains (W59 and W230). It was not possible to assign the resonances to specific residues based on the reported backbone assignment. However, both W59 and W230 are located in helices that form the 14-3-3σ binding groove (Figure 4, panel B). The indole side chain of W230 protrudes into the binding groove and in the crystal structure is observed to form a hydrogen bond with the side chain of the MDM2 peptide H184 residue, which is consistent with the previously reported structure with a mono-phosphorylated MDM2^pS186^ peptide.^17^ The side chain of W59 is proximal to that of V134 for which the corresponding resonance also undergoes CSP – both form part of the 14-3-3σ dimer interface.

In addition to CSPs, the resonance corresponding to E182, which is also located in the binding groove, broadened such that it was not visible in the spectrum of the 14-3-3σ/MDM2 complex. This is also indicative of an interaction between the amino acid residue and the peptide. It is consistent with the crystallography data that shows E182 forms a polar contact with the MDM2 peptide which is again consistent with the previous report.^17^

CSPs were not discernible for the key residues that form the phosphate binding pocket, as observed from the crystal structure (R56, R129, Y130). R129 and Y130 were not assigned in the original backbone assignment of 14-3-3σ,^38^ and so signals for these residues were not identified in our spectra. A small CSP upon phosphopeptide binding for R56 has been reported previously,^39^ but because it lies in a densely populated region of the spectrum it could not be measured from our data.

CSPs were also observed for residues distal to the binding groove: G94, K140, S149, T165, D204 (Figure 4, panels A-B). CSPs for G94 and S149 have been previously reported for 14-3-3σ in complex with a peptide motif mimicking the Tau protein.^39^ These CSPs could reflect conformational changes to 14-3-3σ structure upon peptide binding. T165 and D204 are seen to form hydrogen bond in the crystal structure of the binary complex. Together, the NMR data confirmed that the MDM2_161-191_^pS166/pS186^ peptide was binding in the 14-3-3σ groove.

To probe the dynamics of the 14-3-3σ/MDM2 interaction a titration experiment was conducted. ^1^H-^15^N TROSY spectra of 14-3-3σ were collected in the presence of increasing concentrations of the MDM2_161-191_^pS166/pS186^ peptide. Intensities of resonances corresponding to apo-14-3-3σ decreased as the concentration of peptide was increased. Conversely, resonances corresponding to the 14-3-3σ/MDM2 complex increased as the concentration of peptide was increased, with the exception of that for E182 which broadened such that it could no longer be observed. This was most notable for residues V134, T165, F176, E182, and one of the tryptophan indoles (W59 or W230, Figure 4, panel C). Saturation of 14-3-3σ was observed when one molar equivalent of peptide was introduced relative to the 14-3-3σ monomer (i.e. 100 µM 14-3-3σ and 100 µM MDM2_161-191_^pS166/pS186^ peptide). This provides further evidence for 2:1 monomer-peptide binding stoichiometry, again because a greater excess of peptide would be expected to be required to reach saturation if two peptides were simultaneously engaging a 14-3-3σ dimer. For residues V134, T165 and either W59 or W230, two distinct resonances were observed when 0.5 molar equivalents of peptide were introduced relative to the 14-3-3σ monomer (i.e. 100 µM 14-3-3σ and 50 µM MDM2_161-191_^pS166/pS186^ peptide, Figure 4, panel C). This is intriguing because it indicates that the two states observed in the ^1^H-^15^N TROSY spectra are in slow exchange on the NMR timescale. These two states may reflect a 14-3-3σ conformational change that is induced by MDM2 peptide binding. As only one set of NMR signals (bound and free) are observed for each residue during the titration, symmetry of the dimer is preserved throughout, indicating that the MDM2 peptide binding event itself is in fast exchange between the two available binding grooves present in the 14-3-3 dimer. This complex and dynamic binding mechanism could relate directly to the entropy-driven nature of the interaction as shown by ITC. Furthermore, the conformational change induced upon MDM2 peptide binding could be responsible for the biphasic FP curves as a result of a reduction in fluorophore rotational freedom.

### The 14-3-3σ/MDM2 PPI is Amenable to Small Molecule Stabilisation

Stabilisation of the 14-3-3σ/MDM2 PPI using ‘molecular glues’ (i.e. small molecules that enhance the affinity of a PPI by forming a ternary protein-protein-drug complex) could be an effective strategy for enhancing p53 tumour suppressor activity. To investigate the feasibility of this, the effect of fusicoccin A (FC-A) on the PPI was studied using FP, ITC and native MS. FC-A is a fungal metabolite that predominantly stabilises mode III 14-3-3 PPIs whereby the binary complex creates a ligand binding pocket.^40^ However, FC-A has also been shown to stabilise peptides containing mode I/II type di-phosphorylated binding motifs such as that for CFTR.^27^

FP experiments were conducted to establish if FC-A increased the affinity of the MDM2_161-191_^pS166/pS186^ peptide for 14-3-3σ. 14-3-3σ was titrated to a fixed concentration of the fluorescently labelled MDM2_161-191_^pS166/pS186^ peptide in the presence of increasing concentrations of FC-A (Figure 5, panel A). The effective K_d_ decreased in a FC-A concentration dependent manner from 3.5 ± 2.1 µM in the control experiment to a minimum of 0.60 ± 0.10 µM in the presence of 1.0 mM FC-A. This represented a 5.8-fold stabilisation of the 14-3-3σ/MDM2 PPI. Titration of FC-A to a fixed concentration of 14-3-3σ and the MDM2_161-191_^pS166/pS186^ peptide tracer also showed FC-A dose-dependent stabilisation (EC_50_ = 67.1 ± 41.1 µM).

**Figure 5.**
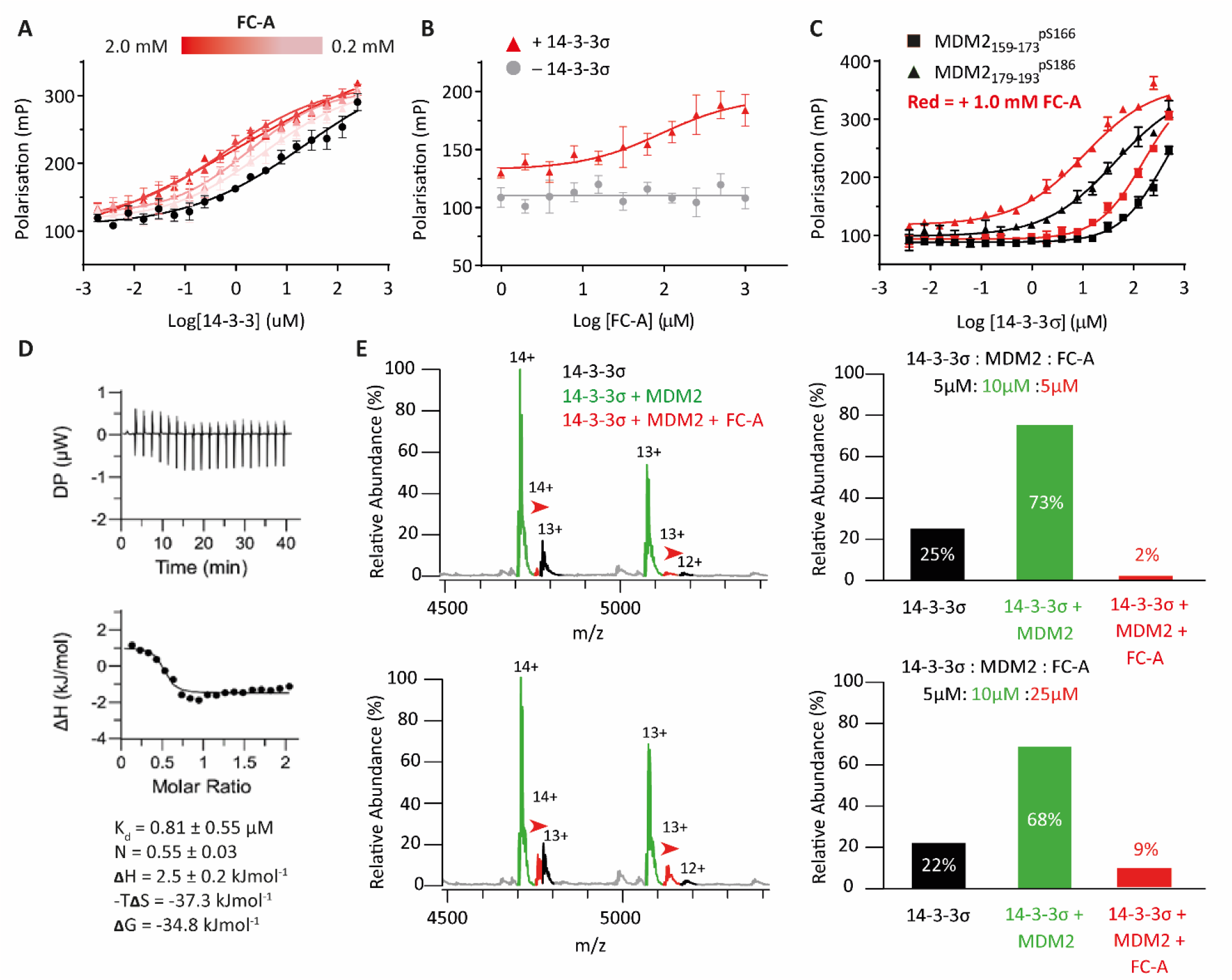
Stabilisation of the 14-3-3/MDM2 PPI by FC-A. **A**: FP data for MDM2_161-191_^pS166/pS186^ peptide binding to 14-3-3σ in the presence of FC-A. 14-3-3σ was titrated to 10 nM FITC-labelled peptide and increasing fixed concentrations of FC-A. **B**: FP dose-response data for FC-A. FC-A was titrated to fixed concentration of 14-3-3σ (1 µM) and FITC-labelled peptide (10 nM). **C**: FP data for MDM2_159-173_^pS166^ and MDM2_179-193_^pS186^ peptide binding to 14-3-3σ. 14-3-3σ was titrated to 10 nM FITC-labelled peptide and 1.0 mM FC-A. All FP experiments were conducted in buffer containing 25 mM HEPES pH 7.5, 100 mM NaCl, 10 mM MgCl2, 0.1% v/v Tween20, 0.1 mg/mL BSA and 1 % v/v DMSO. Error bars represent SD for n=3 replicates. **D**: ITC data for MDM2_161-191_^pS166/pS186^ peptide binding to 14-3-3σ in the presence of FC-A. Acetylated MDM2_161-191_^pS166/pS186^ peptide (1.0 mM) was titrated to 14-3-3σ protein (0.1 mM) in the presence of FC-A (1.0 mM) at 25 °C in buffer containing 25 mM HEPES pH 7.5, 100 mM NaCl, 10 mM MgCl_2_ and 1 % v/v DMSO. E: Native mass spectrometry data showing the FC-A concentration dependent formation of a ternary complex. Red arrows highlight peaks for the ternary complex.

To determine if FC-A had a preferential effect on either phosphorylation site, further 14-3-3σ titrations were carried out using the fluorescently labelled monophosphorylated MDM2_159-173_^pS166^ and MDM2_179-193_^pS186^ peptides. FC-A had a small effect on the 14-3-3σ interaction with the weakly binding MDM2_159-173_^pS166^ peptide whereby the binding curve was shifted slightly to the left, and a higher maximum polarisation value was obtained. As before, saturation was not reached and accurate fitting data were not obtained. FC-A did have a significant effect on the 14-3-3σ interaction with the MDM2_179-193_^pS186^ peptide. FC-A (1.0 mM) stabilised this interaction by 2.5-fold (K_d_ = 24.6 ± 12.0 µM → 10.8 ± 2.9 µM). The maximum polarisation value obtained also increased slightly, likely as a result of conformational restraint of the peptide N-terminus.

ITC was used to further investigate FC-A stabilisation of the 14-3-3σ interaction with the MDM2_161-191_^pS166/pS186^ peptide. The N-terminal acetylated peptide was titrated to a fixed concentration of 14-3-3σ in the presence of FC-A (Figure 5, panel D). In alignment with the previous experiments, the stoichiometry of the interaction was 2:1, 14-3-3 monomer to MDM2 peptide. However, in stark contrast to the binary system the interaction of the MDM2 peptide with 14-3-3σ in the presence of FC-A was an endothermic process entirely dominated by an entropic contribution which could again be explained by a significant hydrophobic effect, or changes in protein structure (Figure 5, panel D). The K_d_ for the interaction in the presence of FC-A could not be accurately measured but did not appear to change significantly, perhaps reflecting the modest stabilising effect: K_d_ = 0.81 ± 0.55 µM, compared to 1.5 ± 0.4 µM for the binary system. Nevertheless, this is a ~2-fold stabilisation that is consistent with the effect seen by FP. Binding was entirely driven by the entropic component which could be a result of a large hydrophobic effect, or of residual structural disorder in the ternary complex.

To confirm that FC-A was engaged in a ternary complex with 14-3-3σ and the MDM2 peptide a series of native MS experiments were conducted. In the presence of two molar equivalents of MDM2 peptide a distribution of 14-3-3σ dimer and 14-3-3σ/MDM2 binary complex was observed (Figure 5, panel E). Upon addition of FC-A a ternary FC-A/MDM2/14-3-3σ complex was formed which comprised a 14-3-3 dimer, a single MDM2 di-phosphorylated peptide, and a single FC-A molecule. As the concentration of FC-A increased the abundance of the ternary complex also increased (Figure 5, panel E and Figure S8). These data confirm that FC-A forms a ternary complex with 14-3-3σ and MDM2 via binding to a single site – the FP data indicate this is most likely the interface of the MDM2 pS186 motif with 14-3-3σ. Furthermore, the data indicate that FC-A modestly promotes complex formation in a concentration-dependent manner.

## Conclusions

Here, we report on a biophysical and structural characterisation of the 14-3-3σ/MDM2 PPI which is an important node in MDM2 and p53 homeostasis and a potential target for therapeutic intervention. The data confirm that affinity of MDM2-derived peptides for 14-3-3σ is dependent on phosphorylation of both S166 and S186 residues which flank a nuclear localisation sequence of MDM2. Peptides mimicking this di-phosphorylated MDM2 motif bind via an entropy-driven process which is unusual in the context of other 14-3-3 PPIs with di-phosphorylated motifs.^34^ This process might be explained by a significant hydrophobic effect, or a change in protein secondary structure upon peptide binding. In contrast to a previous study,^17^ our data show that a single MDM2 peptide occupies both binding grooves present on a 14-3-3σ dimer. Furthermore, the crystallography data confirm that both the pS166 and pS186 engage. However, peptide binding must occur via a fast exchange mechanism on the NMR timescale, indicating that in solution the peptide likely ‘rocks’ between each monomer groove. It highlights the potential significance of cross talk between 14-3-3 monomers in controlling certain PPIs. MDM2 peptide binding also induces a conformational change in the structure of 14-3-3σ which is under a slow exchange mechanism. This is significant because it again highlights the dynamic nature of 14-3-3 proteins. Together the data also point toward cooperativity/negative cooperativity with respect to binding in one versus two monomer binding grooves. The physiological implications of this could be to mask the nuclear localisation sequence on MDM2 leading to cytoplasmic accumulation. However, it is not clear how this interaction promotes MDM2 auto-ubiquitination – this may relate to the aforementioned secondary interaction with the RING domain of MDM2.

Stabilisation of the 14-3-3σ/MDM2 PPI could be an effective strategy for promoting/protecting p53 tumour suppressor activity. Stabilisation of this oncosuppressive interaction may have several advantages over classical MDM2 inhibition. A key issue surrounding inhibition of the MDM2/p53 PPI is an increase in MDM2 protein levels, due to MDM2 being a transcriptional target of p53.^41^ This increase in MDM2 creates a ‘sink’ for inhibitors, which can create a limit to the extent the p53 pathway can be activated. Based on the available biological evidence, stabilisation of the 14-3-3σ/MDM2 may result in MDM2 degradation.^24^ Degradation of MDM2 has recently been shown to indeed dampen the negative feedback mechanism between MDM2 and p53 when compared with MDM2 inhibitors.^42^ Here, we use FP, ITC and native mass spectrometry to demonstrate proof-of-concept for small molecule stabilisation of this PPI using FC-A. Although the stabilising effect of FC-A is modest, the FP data indicate that a ligand binding site is available at the interface of 14-3-3σ and the pS186 motif of MDM2. Thus, the data provide a valuable foundation for further structure-based design efforts to more efficacious and selective molecular glues for this important PPI.

## Author Contributions

JAW conducted the protein expression, peptide synthesis, FP, ITC and NMR experiments in this study. DFK, JB-C and ACL collected and analysed the native MS data. BR-A, HK and PR performed the X-ray protein crystallisation, X-ray diffraction data collection and analysis. PR modelled and refined the MDM2 peptide in the 14-3-3σ groove, deposited the final structure, and wrote the protein crystallography sections in the manuscript. BT and JB assisted with ITC data collection and analysis. SM provided supervision (JAW) and helped to conceive the study. FWM supervised the NMR data collection and analysis, helped to conceive the study, and wrote the manuscript. RD conceived the study, analysed the data and wrote the manuscript. All authors helped to review and edit the final manuscript.

## Acknowledgements

The authors would like to thank Dr. Louise Fairall for assistance with the X-ray diffraction data collection, scaling and analysis. The biophysical and structural research was funded by the University of Leicester and the Engineering and Physical Sciences Research Council (EP/W015803/1). PR was the recipient of a Research Project Prize 2021 (Grant code Plant_EDEM) by the Department of Biological and Agro-alimentary Sciences of the Italian National Research Council. The Eclipse mass spectrometer was funded by the BBSRC (BB/S019456/1). The mass spectrometry research was supported by the Biotechnology and Biological Sciences Research Council (BBSRC) and University of Birmingham funded Midlands Integrative Biosciences Training Partnership (MIBTP2) (BB/M01116X/1).

## Footnotes

§ To accurately calculate peptide concentration the TFA content of the MDM2161-191^pS166/pS186^ peptide was determined by ^19^F-NMR (see Fig. S5 and SI for details).

