## Supporting Information for "Characterising the Protein-Protein Interaction Between MDM2 and 14-3-3σ; Proof of Concept for Small Molecule Stabilisation"

a. Leicester Institute for Structural and Chemical Biology, University of Leicester, University Road, Leicester, LE1 7RH, UK.
b. Mechanisms of Cancer and Aging Laboratory, Department of Molecular and Cell Biology, University of Leicester, University Road, Leicester, LE1 7RH, UK
c. Department of Molecular and Cell Biology, University of Leicester, University Road, Leicester, LE1 7RH, UK
d. School of Biosciences, University of Birmingham, Edgbaston, Birmingham, B15 2TT, UK
e. School of Chemistry, University of Leicester, University Road, Leicester, UK, LE1 7RH, UK.
f. FoodLab, Faculty of Health Sciences, Universitat Oberta de Catalunya, Barcelona, Spain.
g. Josep Carreras Leukaemia Research Institute, Ctra de Can Ruti, Camí de les Escoles, s/n, 08916 Badalona, Barcelona, Spain.
h. Institute of Agricultural Biology and Biotechnology, C.N.R., Unit of Milan, Milano, Italy

### Supporting Information

#### Supporting Figures

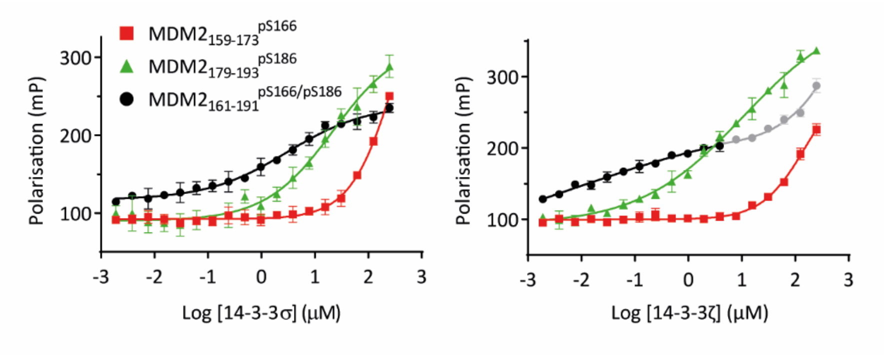

| Peptide | 14-3-3σ $K_d$ (μM) | 14-3-3ζ $K_d$ (μM) |
|---|---|---|
| $MDM2_{159-173}^{pS166}$ | - | - |
| $MDM2_{179-193}^{pS186}$ | 22.3 ± 21.5 | 10.7 ± 24.4 |
| $MDM2_{161-191}^{pS166/pS186}$ | 3.0 ± 1.6 | 0.042 ± 0.02 |

**Figure S1.** Duplicate fluorescence polarisation data in support of Figure 2: data for MDM2 peptide binding to 14-3-3σ (**A**) and 14-3-3ζ (**B**). 14-3-3 protein was titrated to 10 nM FITC-labelled peptide in buffer containing 25 mM HEPES pH 7.5, 100 mM NaCl, 10 mM $MgCl_2$, 0.1% v/v Tween20, 0.1 mg/mL BSA and 1 % v/v DMSO. Error bars represent SD for n=3 replicates.

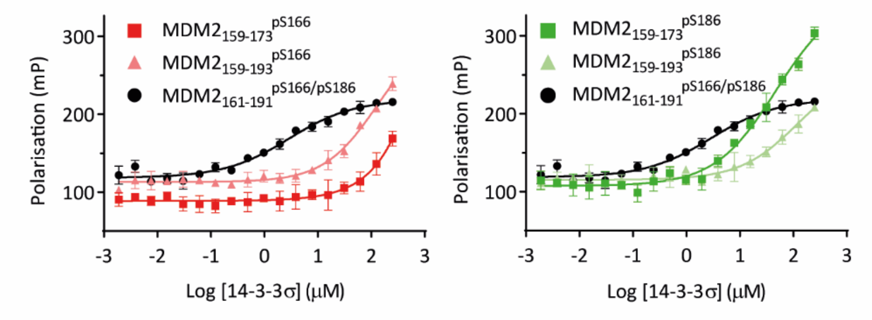

| Peptide | 14-3-3σ $K_d$ (μM) |
|---|---|
| $MDM2_{159-173}^{pS166}$ | - |
| $MDM2_{159-193}^{pS166}$ | - |
| $MDM2_{179-193}^{pS186}$ | 26.2 ± 17.9 |
| $MDM2_{159-193}^{pS186}$ | > 124 |
| $MDM2_{161-191}^{pS166/pS186}$ | 3.0 ± 1.6 |

**Figure S2**. FP data showing that peptide length does not account for the increased affinity of the $MDM2_{161-191}^{pS166/pS186}$ for 14-3-3σ. 14-3-3σ protein was titrated to 10 nM FITC-labelled mono-phosphorylated peptides of different lengths (15mer and 35mer) in buffer containing 25 mM HEPES pH 7.5, 100 mM NaCl, 10 mM $MgCl_2$, 0.1% v/v Tween20, 0.1 mg/mL BSA and 1 % v/v DMSO. Error bars represent SD for n=3 replicates.

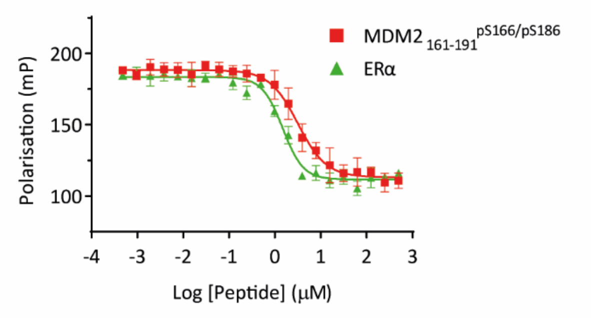

| Peptide | $IC_{50}$ (μM) |
|---|---|
| $MDM2_{161-191}^{pS166/pS186}$ | 3.2 ± 0.5 |
| ERα | 1.5 ± 0.2 |

**Figure S3.** FP competition assay data. $MDM2_{161-191}^{pS166/pS186}$ and ERα peptides were titrated into 10 nM FITC- labelled $MDM2_{161-191}^{pS166/pS186}$ peptide and 10 μM 14-3-3σ in buffer containing 25 mM HEPES pH 7.5, 100 mM NaCl, 10 mM $MgCl_2$, 0.1% v/v Tween20, 0.1 mg/mL BSA and 1 % v/v DMSO. Error bars represent SD for n=3 replicates.

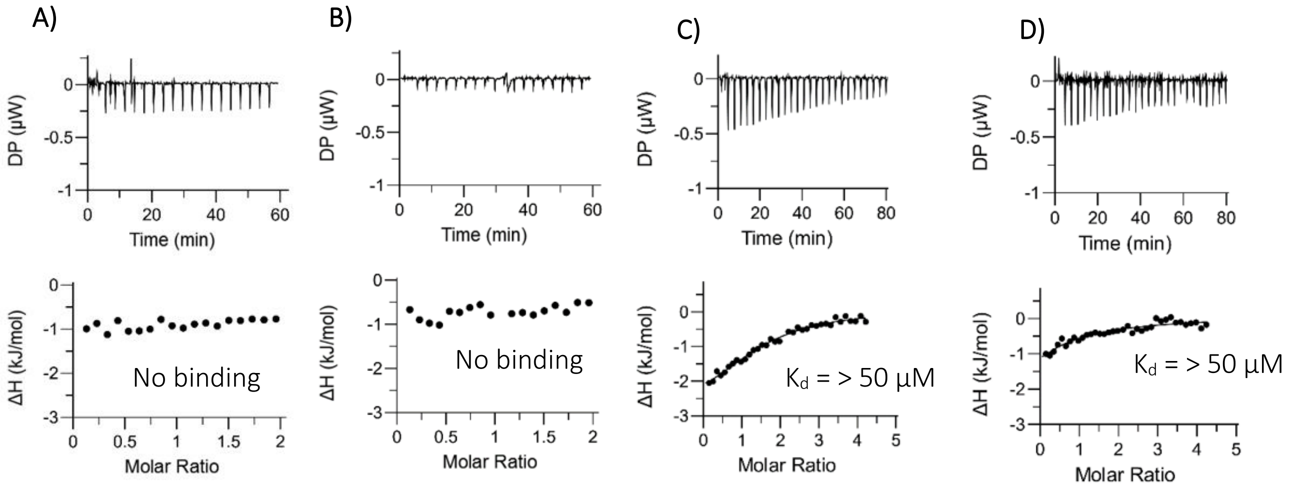

**Figure S4.** ITC data for MDM2 mono-phosphorylated peptides binding to 14-3-3. Acetylated peptides (1.0 mM, syringe) were titrated into 14-3-3 protein (100 μM, cell) at 25 °C in buffer containing 25 mM HEPES pH 7.5, 100 mM NaCl, 10 mM $MgCl_2$ and 1 % v/v DMSO. (**A**) $MDM2_{159-173}^{pS166}$ 14-3-3σ. (**B**) $MDM2_{179-193}^{pS186}$ 14-3-3σ. (**C**) $MDM2_{159-173}^{pS166}$ into 14-3-3ζ. (**D**) $MDM2_{179-193}^{pS186}$ into 14-3-3.

A

B)

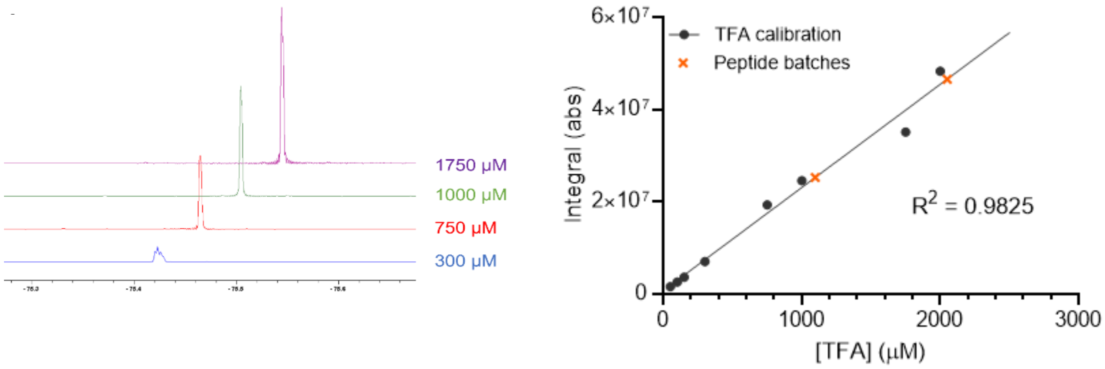

C)

| **TFA:** 114 g/mol<br>**Peptide:** 3,872 g/mol | Ac-MDM2$_{161\text{-}191}$$^{pS166/pS186}$ peptide **Batch 1** | Ac-MDM2$_{161\text{-}191}$$^{pS166/pS186}$ peptide **Batch 2** |
|---|---|---|
| **Integral of 19F resonance at −75 ppm (abs)** | 2.526 x $10^7$ | 4.660 x $10^7$ |
| **Extrapolated concentration (mM)** | 1.1 | 2.1 |
| **Sample volume (µL)** | 600 | |
| **TFA (moles)** | 6.6 x $10^{-7}$ | 12.6 x $10^{-7}$ |
| **TFA (mg)** | 0.08 | 0.14 |
| **Peptide amount in 250 µM stock (mg)** | 0.57 | |
| **TFA (%)** | 14 | 24 |
| **TFA correction factor** | 1.14 | 1.24 |

**Figure S5.** $^{19}$F-NMR experiments to determine % TFA content in MDM2$^{pSer166/pSer186}$ di-phosphorylated 31mer peptide batches. (**A**) $^{19}$F-NMR calibration spectra showing TFA peak at −75 ppm. Peak offset of 0.05 for display purposes. (**B**) TFA content calibration curve. Plot of absolute integral of $^{19}$F-NMR resonance at −75 ppm against TFA concentration (µM). The integrals obtained, and corresponding TFA concentrations, for the peptide batches are shown by an orange cross. (**C**) Table calculating the TFA correction factor applied to each batch of peptide.

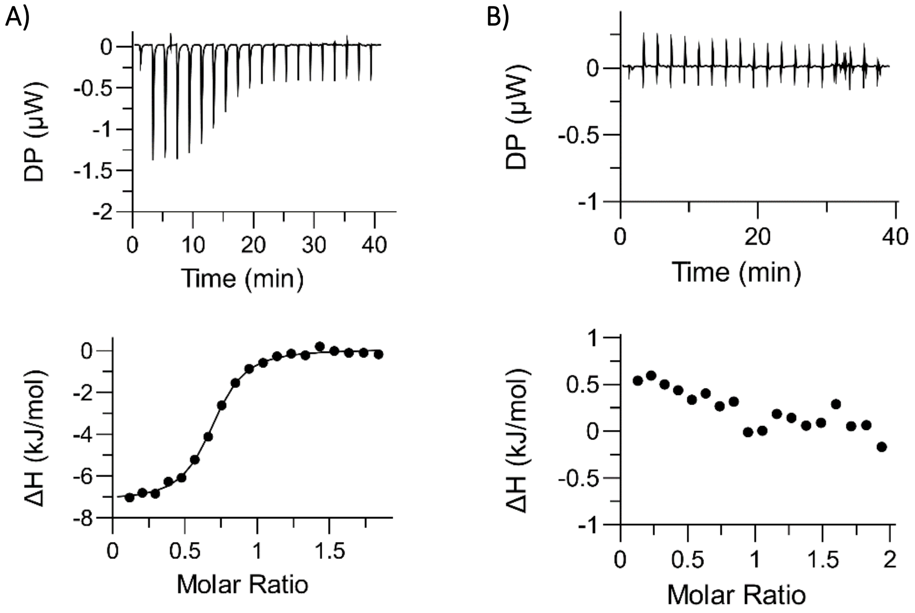

| Parameter | Value |
|---|---|
| N | 0.701 ± 0.043 sites |
| $K_d$ | 2.0 ± 0.2 μM |
| ΔH | -6.3 ± 0.8 μM |
| -TΔS | -26.1 kJ/mol |
| ΔG | -32.6 kJ/mol |

**Figure S6.** ITC data in support of Figure 2. (**A**) Duplicate data for titration of the acetylated MDM2$_{161\text{-}191}^{pS166/pS186}$ peptide (1.0 mM, syringe) into 14-3-3σ protein (100 μM, cell) at 25 °C in buffer containing 25 mM HEPES pH 7.5, 100 mM NaCl, 10 mM $MgCl_2$ and 1 % v/v DMSO. (**B**) Control experiment whereby acetylated MDM2$_{161\text{-}191}^{pS166/pS186}$ peptide (1.0 mM, syringe) was titrated into buffer (cell).

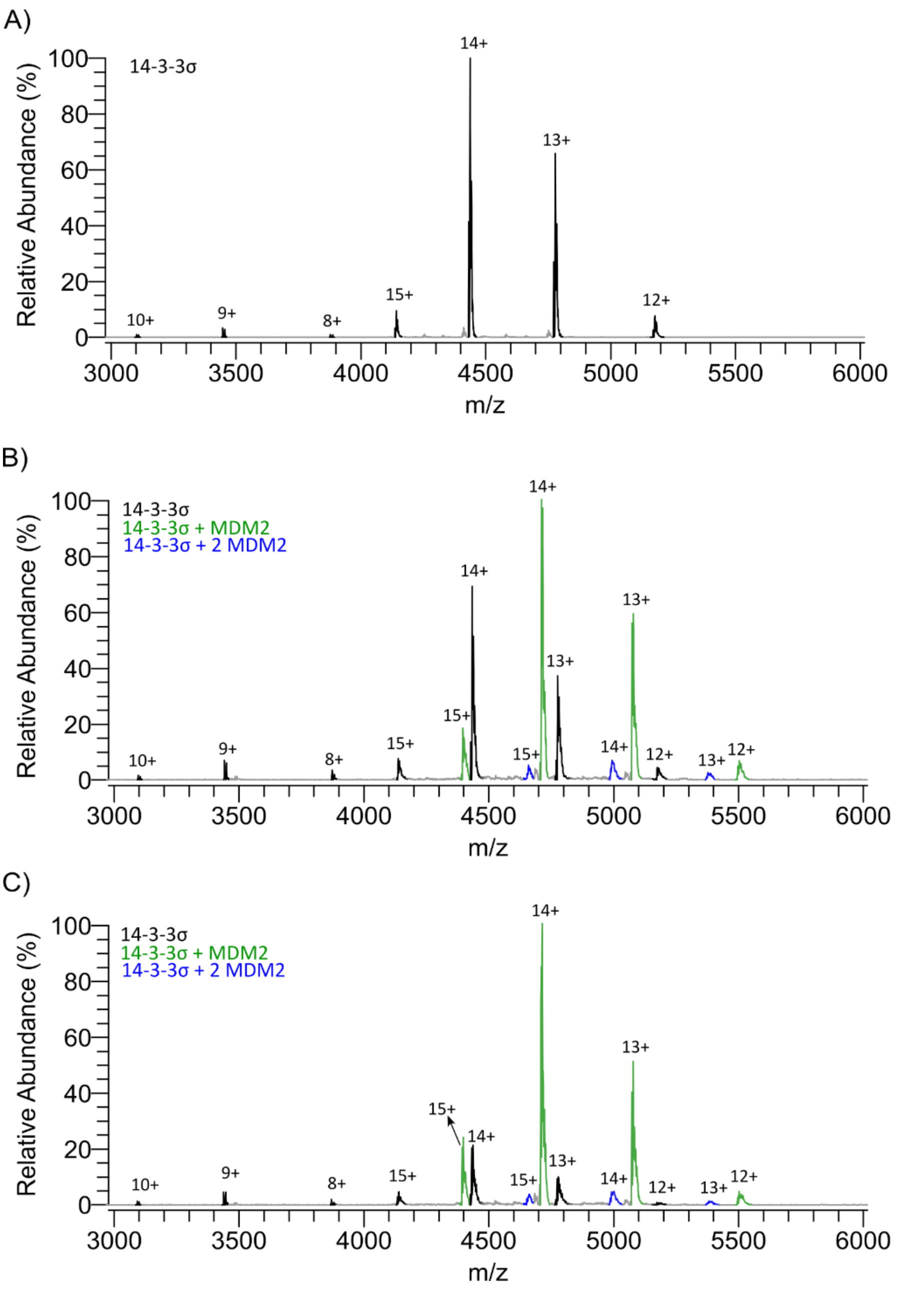

**Figure S7.** Native mass spectrometry data showing binding of 14-3-3 to MDM2. (**A**) Native mass spectrum of 14-3-3σ alone showing 14-3-3σ is dimeric. (**B**) Native mass spectrum of 14-3-3σ incubated with 1 equivalent of MDM2. (**C**) Native mass spectrum of 14-3-3σ incubated with 2 equivalents of MDM2. Peaks corresponding to 14-3-3 σ alone are highlighted in black and the binary complexes with single bound and double bound MDM2 complexes are highlighted in green and blue respectively. Low abundant peaks in 3000-4000 m/z region correspond to the 14-3-3σ monomer.

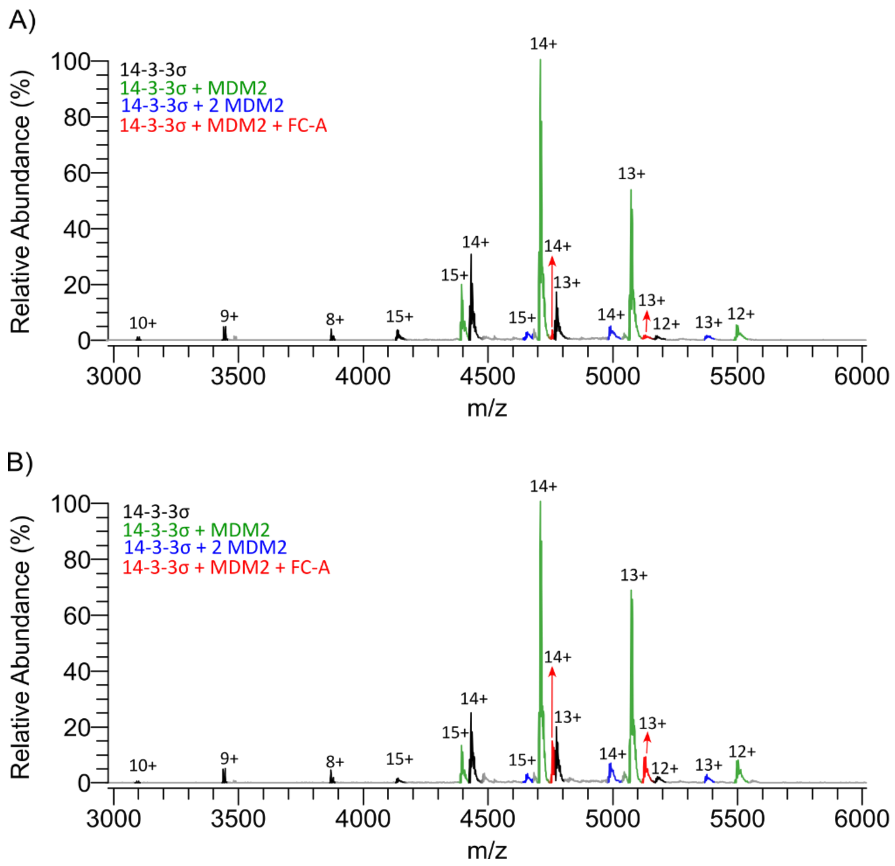

**Figure S8.** Native mass spectrometry data showing ternary complex formation upon binding of FC-A to the 14-3-3/MDM2 binary complex in a concentration-dependent manner. (**A**) 14-3-3 incubated with 2 molar equivalents of MDM2 and 1 molar equivalents of FC-A. (**B**) 14-3-3 incubated with 2 molar equivalents of MDM2 and 5 molar equivalents of FC-A. Peaks corresponding to 14-3-3 σ alone are highlighted in black and the binary complexes with single bound and double bound MDM2 complexes are highlighted in green and blue respectively. The ternary complex is highlighted in red. Low abundant peaks in 3000-4000 m/z region correspond to 14-3-3σ monomer.

| Parameter | Data |
|---|---|
| PDB ID | 8P0D |
| Beamline, date of collection | Diamond I04, 27.11.2021 |
| Structure contents | 14-3-3σ: MDM2$_{161\text{-}191}^{pS166/pS186}$ peptide |
| Wavelength (Å) | 0.91001 |
| Det. Dist., $d_{max}$ (mm, Å) | 1.25 |
| Photon flux ($10^{11}$photons/s) | 1.24 |
| Transmission (%) | 45 |
| Number of images | 3,600 |
| Oscillation range (°) | 0.1 |
| Exposure time (s) | 0.04 |
| Space Group | C 2 2 $2_1$ |
| Cell edges: a, b, c (Å) | 82.59, 111.82 62.55 |
| Resolution Range (Å) | 41.69-1.31 (1.33-1.31) |
| $R_{merge}$ | 0.139 (3.200) |
| $R_{meas}$ | 0.145 (3.324) |
| Observations | 943,346 (46,981) |
| Unique observations | 69,740 (3,449) |
| Average I/$\sigma$(I) | 9.7 (0.5) |
| Completeness | 100.0 (99.7) |
| Multiplicity | 13.5 (13.6) |
| $CC_{1/2}$ | 0.998 (0.299) |

**Table S1.** Crystal X-ray diffraction data collection parameters and data processing statistics for 14-3-3σ: MDM2$_{161\text{-}191}^{pS166/pS186}$ peptide complex. Values in parentheses refer to the highest resolution shell.

| Parameter | Data |
|---|---|
| PDB ID | 8P0D |
| Structure contents | 14-3-3σ: MDM2$_{161\text{-}191}^{pS166/pS186}$ peptide |
| Space Group (Z) | C 2 2 $2_1$ |
| Resolution range (Å) | 18.64-1.31 (1.33-1.31) |
| Reflect.s working set | 69663 |
| Reflect.s free set | 3384(75) |
| $R_{work}/R_{free}$ | 0.1775/0.2036 (0.3468/0.3331) |
| $Rmsd_{bonds}$ (Å) | 0.012 |
| $Rmsd_{angles}$ (°) | 1.17 |
| Ramachandran fav. (%) | 96.6 |
| Ramachandran allow. (%) | 99.3 |
| Ramachandran outliers (%) | 0.7 |
| $\langle B \rangle_{prot}$ ( $\langle B \rangle_{wat}$) (Å$^2$) | 21.37 (38.48) |

**Table S2.** Crystal structures refinement statistics for 14-3-3σ: MDM2$_{161\text{-}191}^{pS166/pS186}$ peptide complex. Values in parentheses refer to the highest resolution shell.

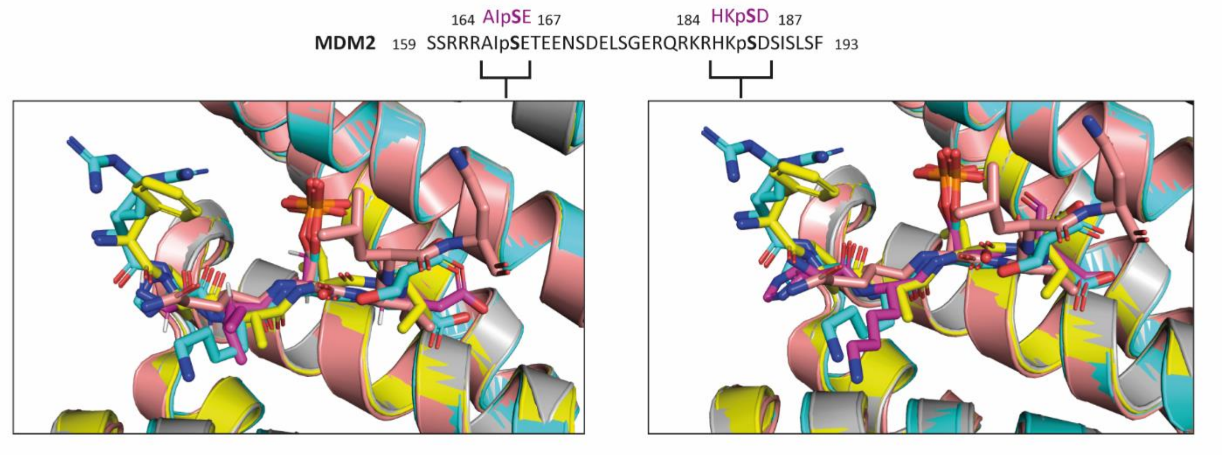

**Figure S9.** Crystal structures showing the two binding stretches of the MDM2 peptide bound in the 14-3-3σ binding groove (14-3-3 shown in grey, peptides shown in magenta, PDB: 8P0D) overlaid with the crystal structures of (1) 14-3-3σ in complex with a peptide mimicking ERα C-terminus (yellow, PDB: 4JC3);[1] (2) 14-3-3σ in complex with a peptide mimicking the MDM2 pS186 binding motif (cyan, PDB: 6YR6);[2] and (3) 14-3-3ζ in complex with a peptide mimicking the CFTR pS768 binding motif.[3]

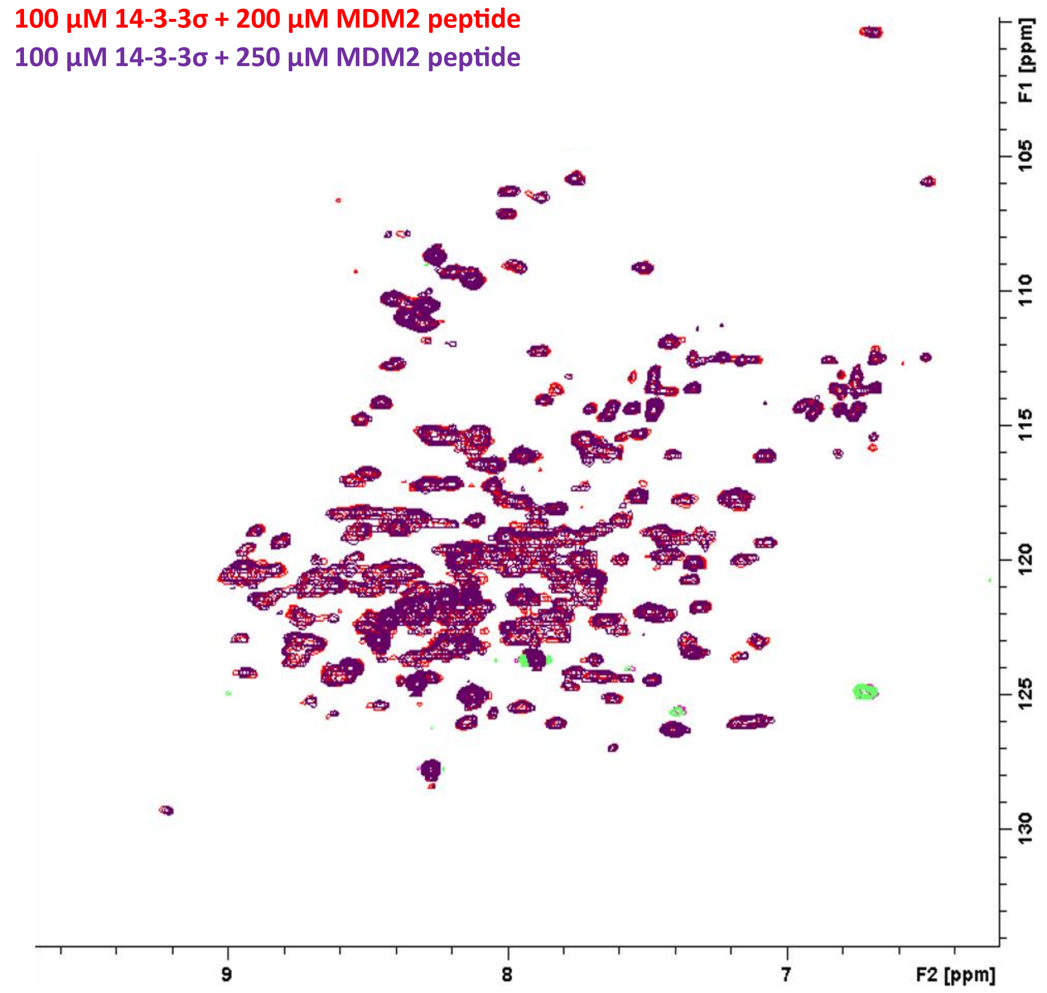

**Figure S10. NMR data in support of Figure 4.** Superimposed $^{1}H$-$^{15}N$ TROSY spectra of $^{15}N$-labelled apo-14-3-3σ (100 µM, black) and $^{15}N$-labelled 14-3-3σ (100 µM) in complex with the $MDM2_{161\text{-}191}^{pS166/pS186}$ peptide (250 µM) (red). All protein concentrations are given as 14-3-3 monomer concentrations.

| Residue | Apo-14-3-3σ | | 14-3-3σ + MDM2 | | CSP | | |
|---|---|---|---|---|---|---|---|
| | δ ($^{15}N$) [ppm] | δ ($^{1}H$) [ppm] | δ ($^{15}N$) [ppm] | δ ($^{1}H$) [ppm] | δ ($^{15}N$) [ppm] | δ ($^{1}H$) [ppm] | **Δδ ($^{1}$H, $^{15}$N) [ppm]** |
| S45 | 108.4 | 7.8 | 109.1 | 8.0 | 0.7 | 0.2 | 0.34 |
| S63 | 114.8 | 8.5 | 114.7 | 8.5 | 0.1 | 0.0 | 0.02 |
| G54 | 109.0 | 7.5 | 108.8 | 7.5 | 0.2 | 0.0 | 0.04 |
| G94 | 106.5 | 8.1 | 106.3 | 8.0 | 0.2 | 0.1 | 0.14 |
| S105 | 114.2 | 8.5 | 114.4 | 8.5 | 0.2 | 0.0 | 0.04 |
| V134 | 107.2 | 6.5 | 105.9 | 6.5 | 1.3 | 0.0 | 0.26 |
| K140 | 119.5 | 6.9 | 119.3 | 7.1 | 0.2 | 0.2 | 0.24 |
| S149 | 112.7 | 8.0 | 112.0 | 7.8 | 0.7 | 0.2 | 0.24 |
| T165 | 100.3 | 6.8 | 100.4 | 6.6 | 0.1 | 0.3 | 0.32 |
| F176 | 123.3 | 9.1 | 122.8 | 9.0 | 0.5 | 0.1 | 0.20 |
| S186 | 112.4 | 8.3 | 112.7 | 8.5 | 0.3 | 0.2 | 0.26 |
| D204 | 116.2 | 7.1 | 115.5 | 7.1 | 0.7 | 0.0 | 0.14 |
| L229 | 123.0 | 7.1 | 122.8 | 7.1 | 0.2 | 0.0 | 0.04 |
| W59/ W230 | 129.5 | 9.3 | 129.2 | 9.2 | 0.3 | 0.1 | 0.06 |

**Table S3.** NMR chemical shift perturbation data in support of Figure 4, panel B. Data for notable combined $^{1}$H and $^{15}$N chemical shift perturbations in the spectrum of $^{15}$N-labelled 14-3-3σ (100 µM) in complex with the $MDM2_{161-191}^{pS166/pS186}$ peptide (200 µM). $^{1}$H, $^{15}$N combined chemical shift changes, Δδ ($^{1}$H, $^{15}$N), upon binding were calculated as $\Delta\delta^{1}H, ^{15}N = |\Delta\delta^{1}H| + |0.2*\Delta\delta^{15}N|$.

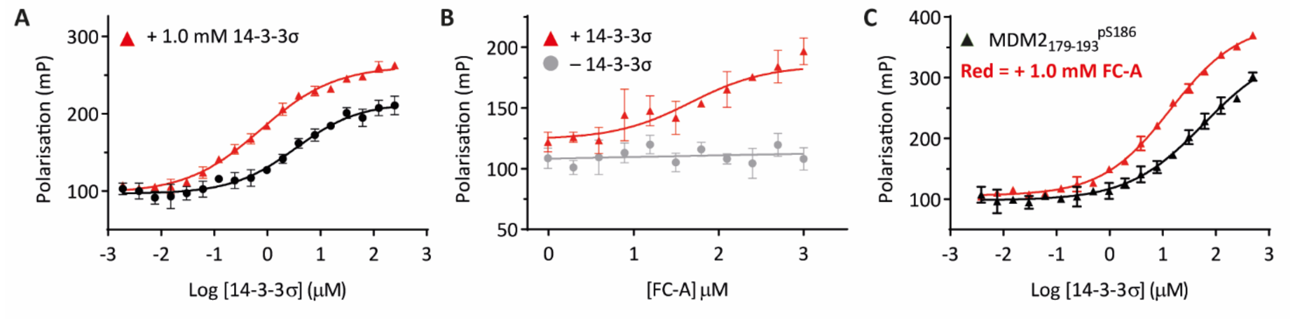

| Experiment | $K_d/EC_{50}$ (µM) |
|---|---|
| A: DMSO Control | 3.4 ± 1.4 |
| A: +FC-A (1.0 mM) | 0.75 ± 0.14 |
| B: FC-A Titration | 47.0 ± 40.1 |
| C: DMSO Control | 59.2 ± 57.7 |
| C: +FC-A (1.0 mM) | 14.1 ± 1.9 |

**Figure S11.** Replicate FP Data in support of Figure 5. **A**: $MDM2_{161-191}^{pS166/pS186}$ peptide binding to 14-3-3σ in the presence of FC-A. 14-3-3σ was titrated to 10 nM FITC-labelled peptide and 1.0 mM FC-A. **B**: FP dose-response data for FC-A. FC-A was titrated to fixed concentration of 14-3-3σ (1 µM) and FITC-labelled peptide (10 nM). **C**: FP data for $MDM2_{179-193}^{pS186}$ peptide binding to 14-3-3σ. 14-3-3σ was titrated to 10 nM FITC-labelled peptide and 1.0 mM FC-A. All FP experiments were conducted in buffer containing 25 mM HEPES pH 7.5, 100 mM NaCl, 10 mM MgCl2, 0.1% v/v Tween20, 0.1 mg/mL BSA and 1 % v/v DMSO. Error bars represent SD for n=3 replicates

### Experimental Procedures

#### Reagents

All reagents were obtained from commercial sources and used without further purification, unless otherwise stated. Fusicoccin A (FC-A) was provided as a gift from Yusuke Higuchi. It was obtained as a metabolite of wildtype *Phomopsis amygdali* and genetically modified *Phomopsis amygdali* Niigata-2 as reported previously.[4]

#### Peptides and Peptide Synthesis

Phosphopeptides were either prepared in-house or were obtained from China Peptides (see Table S2).

| Peptide | Sequence | Source | Purity |
|---|---|---|---|
| $MDM2_{159\text{-}173}^{pS166}$ | AcHN-$^{159}$SSRRRAI(pS)ETEENSD$^{173}$-$NH_2$ | In house | 97.06% |
| $MDM2_{179\text{-}193}^{pS186}$ | AcHN-$^{179}$RQRKRHK(pS)DSISLSF$^{193}$-$NH_2$ | In house | 90.03% |
| FITC $MDM2_{159\text{-}173}^{pS166}$ | FITC-βA-$^{159}$SSRRRAI(pS)ETEENSD$^{173}$-$NH_2$ | In house | 97.18% |
| FITC $MDM2_{179\text{-}193}^{pS186}$ | FITC-βA-$^{179}$RQRKRHK(pS)DSISLSF$^{193}$-$NH_2$ | In house | 94.25% |
| FITC $MDM2_{161\text{-}191}^{pS166/pS186}$ | FITC-βA-RRRAI(pS)ETEENSDELSGERQRKRHK(pS)DSISL-$NH_2$ | In house | 95.68% |
| $MDM2_{161\text{-}191}^{pS166/pS186}$ | AcHN-$^{161}$RRRAI(pS)ETEENSDELSGERQRKRHK(pS)DSISL$^{191}$-$NH_2$ | China Peptides | 96.83% |
| FITC $MDM2_{159\text{-}193}^{pSer166}$ | FITC-Ahx-$^{159}$SSRRRAI(pS)ETEENSDELSGERQRKRHKSDSISLSF$^{193}$-$NH_2$ | China Peptides | 96.72% |
| FITC $MDM2_{159\text{-}193}^{pSer186}$ | FITC-Ahx-$^{159}$SSRRRAISETEENSDELSGERQRKRHK(pS)DSISLSF$^{193}$-$NH_2$ | China Peptides | 96.94% |
| $ER\alpha_{588\text{-}595}^{pThr594}$ | AcHN-$^{588}$AEGFPA(pT)V$^{595}$-COOH | China Peptides | 97.27 % |

AC: acetylation. Ahx: aminohexanoic acid. FITC: fluorescein isothiocyanate.

**Table S4.** Sequences, sources and purity of synthetic peptides used in the study.

**Peptide Synthesis.** In-house peptides were synthesised by Fmoc solid phase peptide synthesis on the Biotage Initiator+ Alstra$^{TM}$ using Rink amide resin (0.55 mmol/g loading). The resin was washed in DMF after each deprotection. Peptide couplings were performed in DMF using 4.0 eq. Fmoc-amino acid, 2.0 eq HCTU and 4.0 eq. DIPEA (2 M solution in NMP) for 5 minutes at 75 °C. The coupling mixture and time were doubled for coupling of arginine resides and any +3 residues N-terminal of an arginine residue. For phosphorylated serine residues a 2.0 eq. of amino acid was used under the same conditions. Fmoc deprotection was carried out at room temperature using 20 % v/v piperidine in DMF for 20 minutes. The peptides were either acetylated or labelled with fluorescein isothiocyanate (FITC) via a β-alanine linker at the N-terminus before deprotection and cleavage from the resin. N-terminal acetylation was performed using 50.0 eq acetic anhydride and 8.0 eq. DIPEA at room temperature for 10 minutes. Fluorescent labelling was performed using 4.0 eq. FITC, 14.0 eq. DIPEA and 1.2 mL NMP at room temperature for 18 hours.
Post-synthesis the resin was washed in diethyl ether and DCM. The peptide was cleaved from the resin using 4 mL TFA:water:ethanedithiol:trispropylsilane in a ratio of 92.5:2.5:2.5:2.5 with shaking for up to 6 hours. The peptide was precipitated into cold diethyl ether and isolated by centrifugation (2,000 rpm, 4°C, 2 minutes). The peptide was washed twice in cold diethyl ether and lyphollised.

**Peptide Purification.** Peptides were purified by reverse phase HPLC using a Thermo Dionex HPLC system with a Luna 5 µm C18(2) preparative column (Phenomenex, 250 x 21.2 mm) with a flow rate of 10.0 ml/min using a linear gradient or isocratic mixture of water and acetonitrile with 0.1 % TFA as additive. Analysis was performed using a Luna Omega 5 µm C18 column (Phenomenex, 250 x 4.6 mm) with a flow rate of 0.5 mL/min and a linear gradient or isocratic mixture of water and acetonitrile with 0.1 % TFA as additive. UV detection wavelength for both analytical and preparative HPLC was 214 nm. Peptides were also analysed by HRMS.

#### Protein Expression and Purification

**FP, ITC, Crystallography.** *N*-terminal $HIS_6$-tagged full-length 14-3-3σ (P31947) and 14-3-3ζ (P63104) proteins were used for ITC, FP and native mass spectrometry experiments. A $HIS_6$-tagged 14-3-3σ ΔC17 construct was used for protein crystallography. Both proteins were prepared via the following procedure:

$HIS_6$-tagged full-length 14-3-3σ and $HIS_6$-tagged 14-3-3σ ΔC17 were expressed in BL21 (DE3) competent cells with a pPROEX HTb plasmid. A single transformed colony was selected and used to inoculate 20 mL terrific broth (containing 100 mg/L ampicillin) and was grown overnight at 37 °C, 180 rpm. The starter culture was used to inoculate 1 L of terrific broth media (containing 100 mg/L ampicillin) and supplemented with 0.4% v/v glycerol and 0.5 mM $MgCl_2$. The cells were grown at 37 °C, 180 rpm until the $OD_{600}$ reached 0.6 – 0.8. Expression was induced by the addition of 0.4 mM IPTG. Incubation was continued overnight at 25 °C, 180 rpm.

Cells were harvested by centrifugation (5,000 rpm, 4 °C, 20 mins) and resuspended in buffer 50 mM HEPES pH 8.0, 300 mM NaCl, 12.5 mM Imidazole supplemented with DNAase and a Pierce™ protease inhibitor tablet. The cells were lysed by sonication and the addition of lysozyme. The lysate was cleared by centrifugation (13,000 rpm, 4 °C, 50 mins). The clear lysate was loaded onto a $Ni^{2+}$-affinity chromatography column equilibrated with 50 mM HEPES pH 8.0, 300 mM NaCl and 12.5 mM Imidazole. The $Ni^{2+}$-affinity chromatography column was washed with 50 mM HEPES pH 8.0, 300 mM NaCl and 25 mM Imidazole. The protein was eluted with 50 mM HEPES pH 8.0, 300 mM NaCl and 250 mM Imidazole. The proteins were dialysed against buffer containing 25 mM HEPES pH 7.5, 100 mM NaCl and 10 mM MgCl2 and concentrated using a 10kDa cut-off centrifugal filter unit (Merck Millipore). All proteins were stored at -80 °C. Typical protein yields were 100 mg/ L.

The $His_6$-tag was cleaved from the 14-3-3σ ΔC17 by TEV protease overnight at room temperature in buffer containing 25 mM HEPES pH 7.5, 100 mM NaCl, 10 mM $MgCl_2$ and 5 mM β-mercaptoethanol. The molar ratio of 14-3-3σ ΔC17 to TEV protease was 1:20. The $His_6$-cleaved 14-3-3σ ΔC17 was further purified by $Ni^{2+}$-affinity chromatography followed by size-exclusion chromatography using a Superdex 75 30/100 column. The pure protein was concentrated and stored as detailed above.

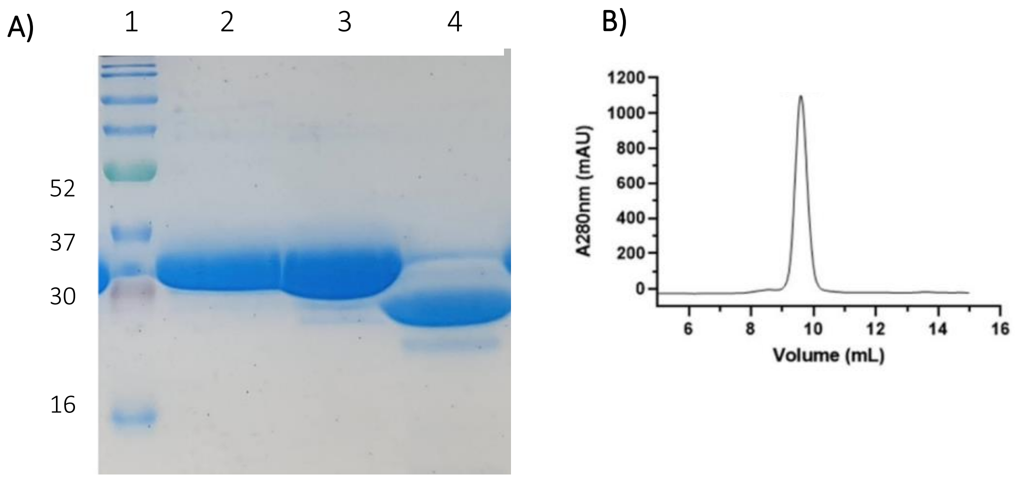

**Figure S12:** (A) SDS-PAGE gel of 14-3-3 proteins used. The Prime-Step™ Prestained Broad Range Protein Ladder from BioLegend (1), 14-3-3ζ (2), 14-3-3σ (3) and 14-3-3σ ΔC (4). (B) Size-exclusion chromatography trace of 14-3-3σ ΔC using a Superdex 75 30/100 column.

**NMR.** $^{15}$N-labelled $HIS_6$-tagged full-length 14-3-3σ was expressed in BL21 (DE3) competent cells with a pPROEX HTb plasmid. A single transformed colony was selected and used to inoculate 50 mL lysogeny broth (containing 100 mg/L ampicillin) and was grown overnight at 37 °C, 180 rpm. A 50 mL starter culture in lysogeny broth (containing 100 mg/L ampicillin) was grown overnight at 37°C, 180 rpm. The cells were sedimented by centrifugation (1,800g for 12 mins at room temperature) and resuspended in 4 mL Modified Spizien's media using $^{15}$N-$NH_4Cl$ as the nitrogen source. The starter culture was used to inoculate 1 L of modified Spizien's minimal media (using $^{15}$N-$NH_4Cl$ as the nitrogen source) supplemented with 100 mg/L ampicillin. The cells were grown at 37 °C, 180 rpm until the $OD_{600}$ reached 0.9-1.0. Expression was induced by the addition of 0.4 mM IPTG. Incubation was continued overnight at 25 °C, 250 rpm. Cells were harvested by centrifugation (5,000 rpm, 4 °C, 20 mins). The protein was purified and the $HIS_6$-tag was removed by TEV protease as detailed above. The protein was stored at -80 °C in buffer containing 25 mM HEPES pH 7.5, 100 mM NaCl, 10 mM $MgCl_2$ and 3 mM $NaN_3$. Typical protein yields were 10 mg/L of culture.

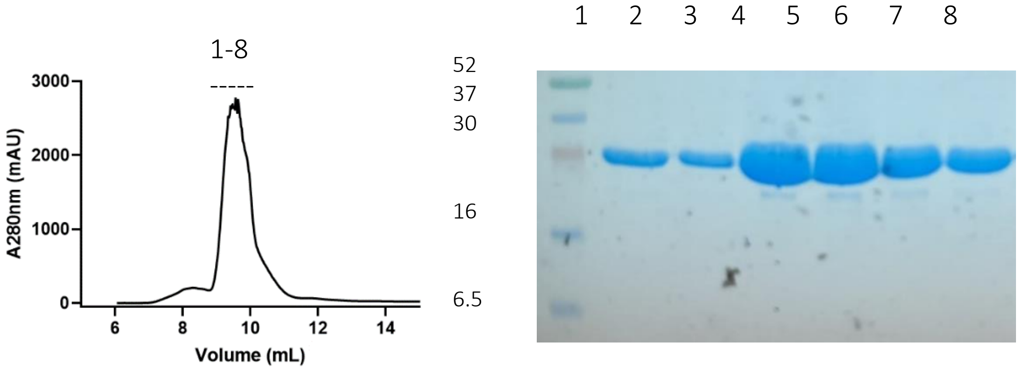

**Figure S13:** SDS-PAGE gel and size-exclusion chromatography trace of $^{15}$N-labelled 14-3-3σ used in NMR experiments. Numbers indicate fractions from size-exclusion chromatography using a Superdex 75 30/100 column. The Prime-Step™ pre-stained Broad Range Protein Ladder from BioLegend was used.

#### Fluorescence Polarisation

Fluorescence polarisation (FP) experiments were conducted at room temperature in buffer containing: 25 mM HEPES pH 7.5, 100 mM NaCl, 10 mM MgCl2, 0.1% v/v Tween20, 0.1 mg/mL BSA and 1 % v/v DMSO using Corning black, round-bottom, low-binding 384-well plates. For all experiments a fixed concentration of 10 nM fluorescently labelled peptide (FITC) was used. All plates were incubated for 10 minutes and shaken for 10 seconds before polarisation was measured using a Hidex Sense Microplate reader with an excitation wavelength of λex: 490 / 20 nm; and an emission wavelength of λem: 535 / 20 nm; mirror dichoric 560; flashes: 20; PMT voltage 750; Z-position: calculated from well. All data was analysed using GraphPad Prism 7 and sigmoidal curves were fitted using the following equation:

Y=Bottom + (Top-Bottom)/(1+10^((Log app.Kd-X)*HillSlope))

Y = mP value
X = log 14-3-3 concentration
Top and bottom = plateaus in mP

#### Isothermal Titration Calorimetry

Isothermal titration calorimetry (ITC) experiments were conducted using the Malvern MicroCal ITC200 instrument (monophosphorylated peptides) or Malvern PEAQ ITC instrument (diphosphorylated peptide). The ITC conditions used are detailed below in Table S3. All experiments were conducted in buffer containing 25 mM HEPES pH 7.5, 100 mM NaCl, 10 mM $MgCl_2$ and 1 % v/v DMSO. Where 2 titrations were conducted the data was merged using Concat32 software (Malvern Instruments Ltd.) All data was analysed using MicroCal PEQ-ITC Analysis Software (single site binding model) to obtain parameters (where applicable).

| Condition | Malvern MicroCal ITC200 | Malvern PEAQ ITC |
|---|---|---|
| Temperature | 25 °C | |
| Stirring speed | 750 rpm | |
| Reference power (DP) | 5 µcal/sec | 10 µcal/sec |
| Feedback | High | |
| Injection spacing | 180 seconds | 120 seconds |
| Initial delay | 60 seconds | |
| Number of injections | 18 | 20 |
| Injection volume | 2 µL* | |

*First injection was 0.4 µL and excluded prior to analysis.

**Table S5.** ITC experimental conditions.

#### Native Mass Spectrometry

Analytical grade ammonium acetate and LC-MS grade Dimethyl Sulfoxide (DMSO) were purchased from Fisher Scientific. 14-3-3 σ was buffer exchanged into 100 mM ammonium acetate pH 6.8 using a 30 kDa molecular weight cut-off Amicon Ultra centrifugal filter (Merck Millipore) and stored at -80 °C prior to use. The lyophilised $MDM2_{161-191}^{pS166/pS186}$ 31mer peptide (see Table S2) was diluted into 100mM ammonium acetate pH 6.8. To form the 14-3-3/MDM2 complex, 14-3-3σ (5 µM) was mixed with 5 µM and 10 µM MDM2 31mer diphosphorylated peptide. To form the stabilised 14-3-3/MDM2 complex, 14-3-3σ (5 µM) was incubated with both a 2-fold excess of the MDM2 31mer diphosphorylated peptide (10 µM) and FC-A (both 5 µM and 25 µM) in 100 mM ammonium acetate pH 6.8 and directly infused into the mass spectrometer. A final concentration of 1.25 % DMSO was used for all experiments.

All native MS experiments were performed on an Orbitrap Eclipse Tribrid mass spectrometer (Thermo Fisher Scientific, Bremen) coupled to a nanoelectrospray source that used gold-coated borosilicate glass capillaries, pulled in-house. Positive ionization mode was used throughout with the capillary voltage set to 1.2 kV. The source temperature was set at 275 °C, in-source dissociation at 25, S-lens RF at 100. High pressure mode was used and a mass range of 2000-8000 *m/z* used to monitor the binding equilibria. Mass spectra were acquired using a maximum ion injection time of 50 ms. The automatic gain control was set to $1x10^6$ and the ions detected in the Orbitrap with resolution set to 7,500. All data was analyzed using XCalibur (*v*.4.1). All proteins and protein complexes were identified based off their theoretical mass.

To quantify each complex, the relative abundance of the complexes observed was assumed to reflect their abundance in solution. Due to overlapping peaks, 15+ ions were excluded from analysis. The quantified complexes included in all calculations were 14-3-3σ, 14-3-3σ/MDM2, 14-3-3σ/FC-A and 14-3-3σ/MDM2/FC-A whereby the relative abundance of each complex was calculated as a percentage of the sum of all other complexes.

#### Protein X-Ray Crystallography

For crystallisation, 10 mg/mL of 14-3-3σ ΔC17 was mixed with MDM2 peptide in a 1:1 molar ratio in 25 mM HEPES pH 7.5, 100 mM NaCl, 10 mM $MgCl_2$ and incubated overnight at 4°C. Vapour diffusion crystallisation 200 nL sitting drops were set up using a Mosquito crystallisation robot (SPT Labtech), mixing the 14-3-3σ ΔC17/MDM2 peptide complex in mother liquor in the following volume ratios: 1:1 (drop 1) and 1:2 (drop 2), using a bespoke crystallography screen of pH and PEG concentrations with 1 M HEPES and 0.19 M $CaCl_2$. All plates were stored refrigerated (4°C). Crystals grew in drops 1 and 2 within a week at the following conditions: 1 M HEPES pH 7.4 to 7.6, 0.19 M $CaCl_2$, 30 to 31 % PEG 400, 5 % glycerol. After 1 month, crystals were fished at 4°C, flash cooled in liquid nitrogen and exposed to X-rays.

For structure determination, the CCP4[5] software package was used with PDB: 3IQJ[6] serving as a template of the 14-3-3σ structure for molecular replacement using PHASER.[7] Further rounds of manual model building and refinement were performed using COOT[8] and REFMAC[9], respectively. Data collection and refinement statistics are shown in Tables S1 and S2.

#### NMR

All spectra were recorded from 100 µM samples of 14-3-3σ in buffer containing 25 mM HEPES, pH 7.5, 100 mM NaCl, 10 mM $MgCl_2$, 3 mM sodium azide and 5% (v/v) $D_2O$ in a 5 mm Shigemi tube (sample volume: 350 µL) using a Bruker 600 MHz AVIII spectrometer operating at 303K. The $^1H^{15}N$ TROSY spectra were recorded with acquisition times of 60 msec in the directly detected dimension and 40 msec on the indirect dimension, with 48 scans per increment (total acquisition time of ~ 5 hours). Spectra were collected for 14-3-3σ only, and in the presence of different molar equivalence of MDM2 peptide (0.25, 0.5, 0.75, 1.0, 2.0 and 2.5 eq). Spectra were processed with Topspin 4.0.6 (Bruker Biospin) to produce the $^1H^{15}N$ TROSY spectra. $^1H$, $^{15}N$ combined chemical shift changes upon binding were calculated as $\Delta\delta^1H, ^{15}N = |\Delta\delta^1H| + |0.2*\Delta\delta^{15}N|$, with Δ the difference between chemical shifts values δ between the apo-14-3-3σ resonances and those in the presence of 2.0 eq of MDM2 peptide.

#### Characterisation of Peptides Synthesised In-House

**Peptide:** MDM2$_{159\text{-}173}^{pS166}$

**Sequence:** AcHN-$^{159}$SSRRRAI(pS)ETEENSD$^{173}$-$NH_2$

**Molecular weight:** 1857.81 g/mol

**ESI-MS m/z:** [M+H] 1858.8094, [M+2H]2+ 929.4108, [M+3H]3+ 619.9421 (see spectra below)

**Purification method:** Linear gradient of 5-95% MeCN in water (0.1 % TFA as additive) over 30 minutes

**HPLC Retention Time:** 18.337 minutes (linear gradient of 5-95% MeCN in water (0.1 % TFA as additive) over 30 minutes) (see spectra below)

**HPLC Purity:** 97.06 % (see spectra below)

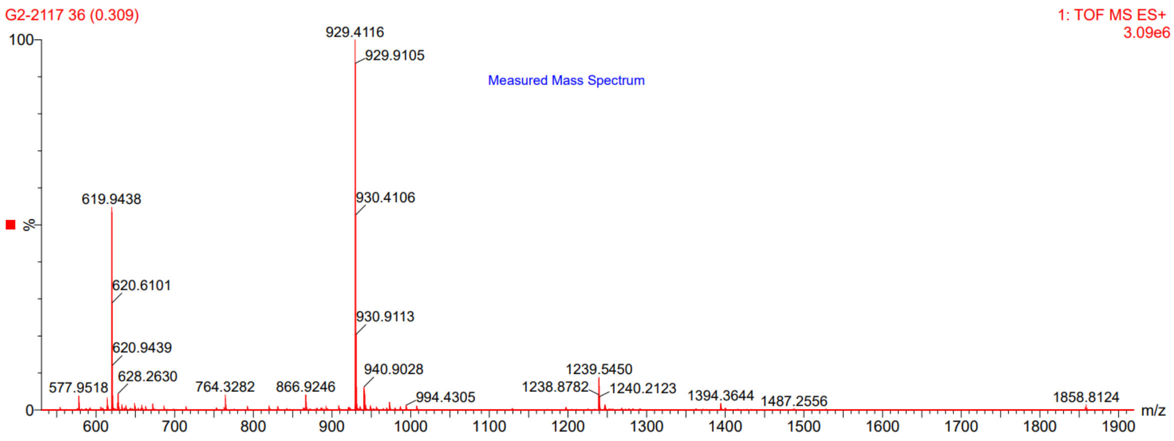

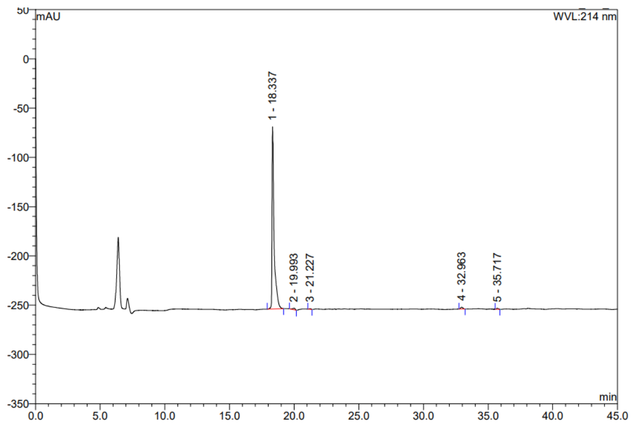

**Peptide:** MDM2$_{179\text{-}193}$$^{pS186}$

**Sequence:** AcHN-$^{179}$RQRKRHK(pS)DSISLSF$^{193}$-$NH_2$

**Molecular weight:** 1966.13 g/mol

**ESI-MS m/z:** [M+H] 1967.0167, [M+2H]2+ 983.5124, [M+3H]3+ 656.3446, [M+4H]4+ 492.5101 (see spectra below)

**Purification method:** Isocratic 22 % MeCN in water (0.1 % TFA as additive) over 30 minutes

**HPLC Retention Time:** 18.930 minutes (linear gradient of 5-95% MeCN in water (0.1 % TFA as additive) over 30 minutes) (see spectra below)

**HPLC Purity:** 90.03 % (see spectra below)

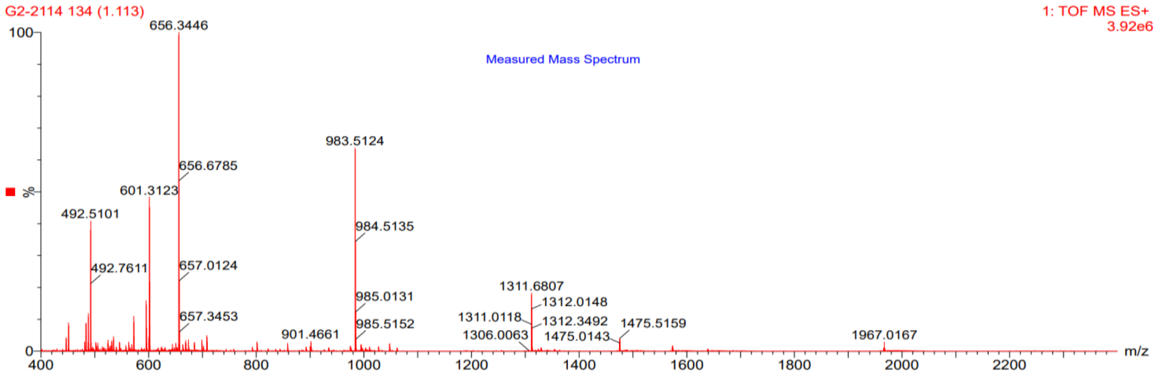

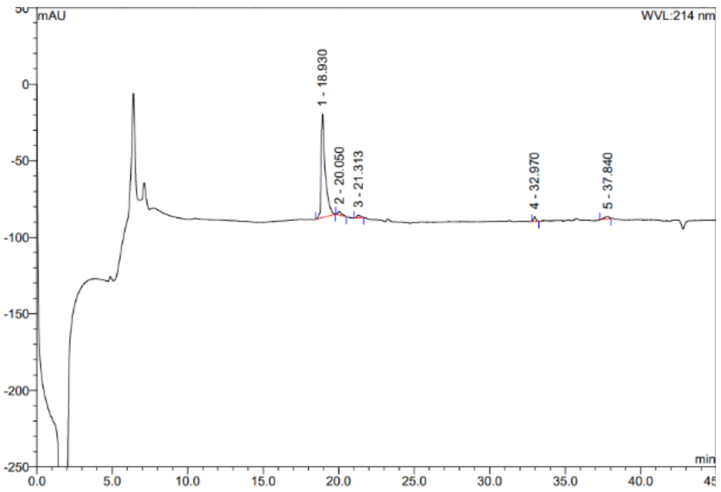

**Peptide:** FITC MDM2$_{159\text{-}173}^{pS166}$

**Sequence:** FITC-βA-$^{159}$SSRRRAI(pS)ETEENSD$^{173}$-$NH_2$

**Molecular weight:** 2276.23 g/mol

**ESI-MS m/z:** [M+3H]3+ 759.2586, [M+4H]4+ 569.9822 (see spectra below)

**Purification method:** Linear gradient of 5-95% MeCN in water (0.1 % TFA as additive) over 30 minutes

**HPLC Retention Time:** 21.130 minutes (linear gradient of 5-95% MeCN in water (0.1 % TFA as additive) over 30 minutes) (see spectra below)

**HPLC Purity:** 97.18 % (see spectra below)

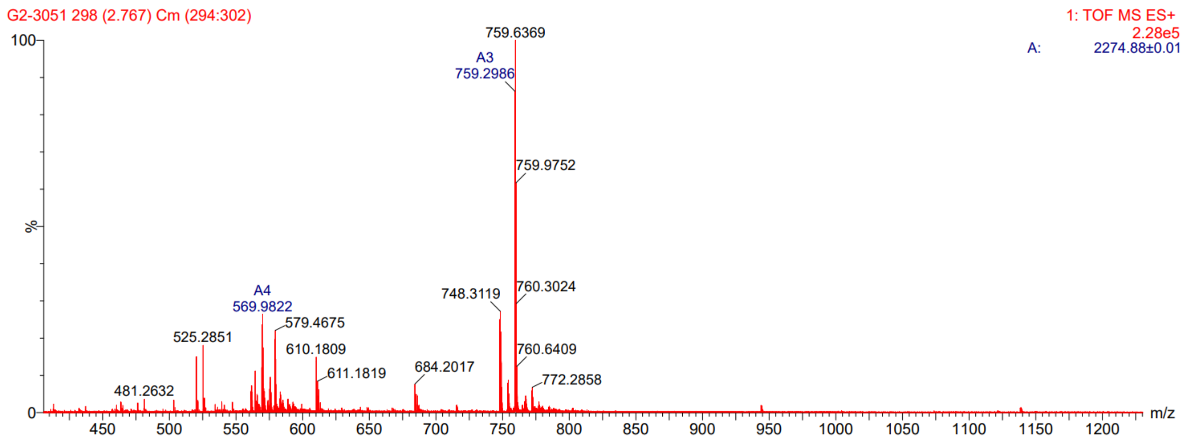

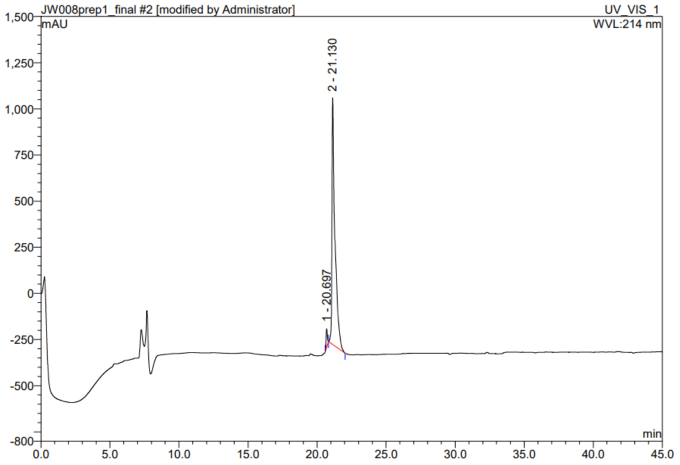

**Peptide:** FITC $MDM2_{179-193}^{pS186}$

**Sequence:** FITC-βA-$^{179}$RQRKRHK(pS)DSISLSF$^{193}$-$NH_2$

**Molecular weight:** 2384.56 g/mol

**ESI-MS m/z:** [M+2H]2+ 1192.5408, [M+3H]3+ 795.3719, [M+4H]4+ 596.7833, [M+5H]5+ 477.6266 (see spectra below)

**Purification method:** Linear gradient of 27-35% MeCN in water (0.1 % TFA as additive) over 30 minutes

**HPLC Retention Time:** 21.707 minutes (linear gradient of 5-95% MeCN in water (0.1 % TFA as additive) over 30 minutes) (see spectra below)

**HPLC Purity:** 94.25 % (see spectra below)

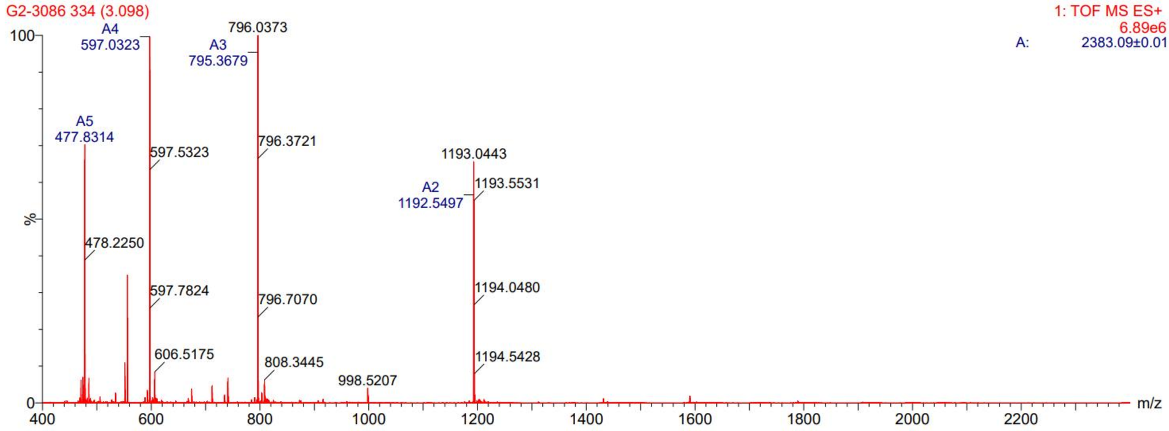

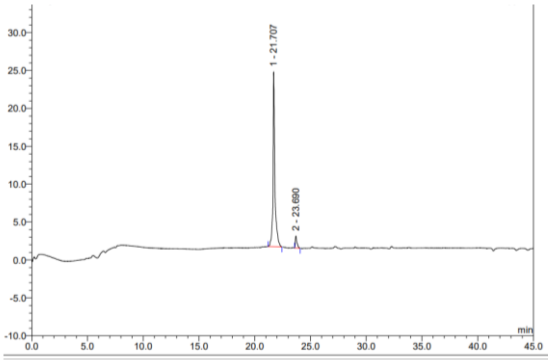

**Peptide:** FITC MDM2$_{161\text{-}191}^{pS166/pS186}$

**Sequence:** FITC-βA-RRRAI(pS)ETEENSDELSGERQRKRHK(pS)DSISL-$NH_2$

**Molecular weight:** 4288.87 g/mol

**ESI-MS m/z:** [M+3H]3+ 1430.3485, [M+4H]4+ 1073.0154, [M+5H]5+ 858.8174, [M+6H]6+ 715.8551, [M+7H]7+ 613.5918 (see spectra below)

**Purification method:** Isocratic 24 % MeCN in water (0.1 % TFA as additive) over 30 minutes

**HPLC Retention Time:** 20.903 minutes (linear gradient of 5-95% MeCN in water (0.1 % TFA as additive) over 30 minutes) (see spectra below)

**HPLC Purity:** 95.68 % (see spectra below)

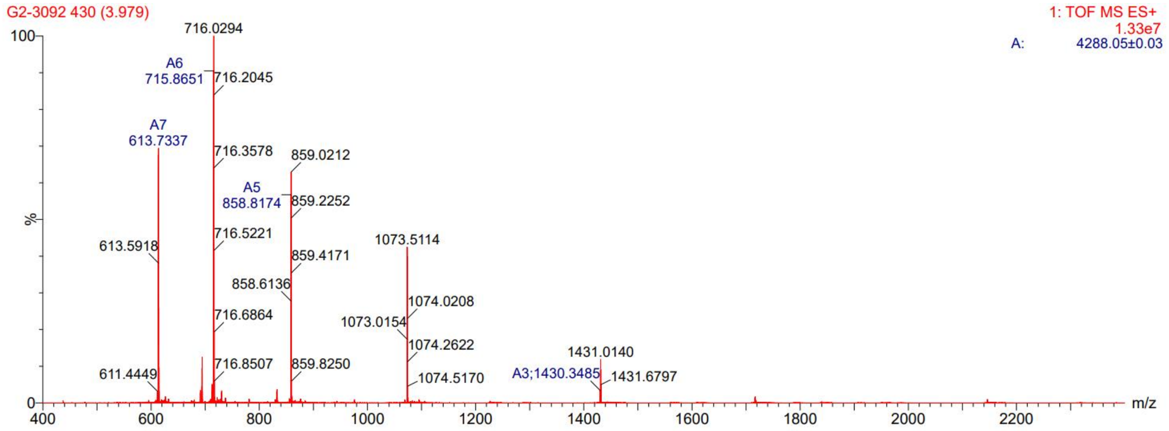

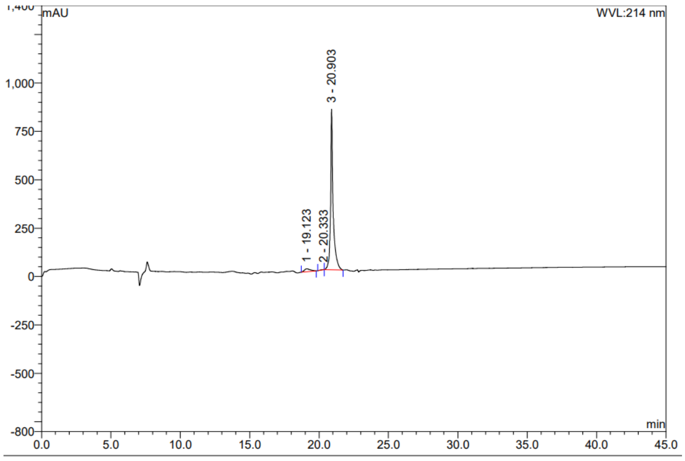
